# A tripartite chemogenetic fluorescent reporter: design and evaluation for visualizing ternary protein complexes

**DOI:** 10.1101/2023.10.19.563144

**Authors:** Sara Bottone, Fanny Broch, Antoine Gedeon, Aurélien Brion, Lina El Hajji, Hela Benaissa, Arnaud Gautier

**Affiliations:** Sorbonne Université, École Normale Supérieure, Université PSL, CNRS, Chimie Physique et Chimie du Vivant (CPCV), 75005 Paris, France; Institut Universitaire de France, Paris, France

## Abstract

Most cellular processes are carried out by multiprotein assemblies. Although various molecular tools exist to visualize binary protein interactions in live cells, the visualization of multiprotein complexes remains a challenge. Here, we report the engineering of a complementation-based approach allowing one to visualize the interaction of three proteins through effective proximity-induced complementation of three fragments of pFAST, a chemogenetic fluorescent reporter that binds and stabilizes the fluorescent state of fluorogenic chromophores (so-called fluorogens). This tripartite-split-pFAST allowed the observation of dynamic ternary protein complexes in the cytosol, at the plasma membrane, in the nucleus and at the junction of multiple organelles, opening great prospects to study the role and function of multiprotein complexes in live cells and in various biologically relevant contexts.

## INTRODUCTION

Physical closeness plays essential roles in biology. Interactions between proteins regulate nearly every major process in cells, from signal transduction to gene regulation, protein transport or metabolism. Close proximity between organelles allows communication through membrane contacts and plays important roles in maintaining cellular homeostasis^1,2^. Various methods have been developed to visualize interactions or proximity in live cells by fluorescence microscopy^3^. Most of them rely either on Förster Resonance Energy Transfer (FRET) or bimolecular fluorescence complementation (BiFC). FRET is a physical effect between two fluorophores in close proximity. As the efficiency of the energy transfer is sensitive to the distance between the two fluorophores, FRET can be used to visualize the proximity of two entities coupled to the appropriate fluorophores^4^. BiFC, on the other hand, is based on the fusion of two non-fluorescent fragments from a split fluorescent protein to two interaction partners^5–7^. Close proximity brings the two non-fluorescent fragments together, allowing the reconstitution of a functional fluorescent protein. BiFC is often preferred as it allows the direct visualization of interactions or proximity with high contrast and its implementation is technically simpler. However, the slow and irreversible complementation of the split fluorescent proteins prevents their general use for the study of dynamic interactions. Split fluorescent reporters with rapid and reversible complementation have been recently developed^8–11^, relying on Fluorescence-Activating and absorption-Shifting Tags (FASTs), small proteins able to bind and stabilize the fluorescent state of synthetic fluorogenic chromophores (so-called fluorogens) that are otherwise dark when free in solution or cells^12–14^. Two non-fluorescent fragments of FASTs are fused to two interaction partners. When in close proximity, a functional split-FAST is reconstituted, enabling fluorogen binding and the formation of a fluorescent assembly. As the binding of the fluorogen is rapid, non-covalent and dynamic, the complementation of split-FASTs is rapid and reversible, allowing the visualization of dynamic protein-protein interactions in real-time^8,11^. Split-FAST has been recently used for developing fluorescent probes enabling to characterize the dynamics of organelle-organelle contact sites^15^, and lipid droplet-organelle interactions^16^.

Although FRET and BiFC have been widely used to study binary protein interactions, the visualization of multiple protein complexes in living cells remains a challenge. Because most cellular processes are carried out by assemblies of more than two proteins^17^, the detection of ternary or even higher order interactions is essential to study and understand the processes that govern cells. Existing tools to analyze three-partner interactions are limited to three-chromophore FRET (3-FRET)^18^, which uses three mutually dependent energy transfer processes between cyan, yellow and red fluorescent proteins, and BiFC-based FRET^19^, which relies on FRET between a cyan fluorescent protein and a split yellow fluorescent protein.

Here, we present the development of a tripartite-split-FAST able to complement when three proteins are proximal or interact (Fig. 1). While various bipartite split reporters exist, the design of tripartite split systems capable of reconstituting when three entities are in close proximity is more challenging, because of the inherent difficulty of reassembling a functional system from three complementary split fragments. Up to now, only the superfolder green fluorescent protein (sfGFP)^20^ and Nanoluciferase^21^ have been split into three fragments. These tripartite systems have been however only used to detect binary interactions with minimal self-assembly background. To develop a tripartite-split-FAST allowing the visualization of ternary interactions in living cells, we first identified a second split site able to generate alternative functional bipartite-split-FAST variants (Fig. 1). The combination of the two functional split sites allowed us to generate three complementary fragments that only assemble when in proximity (Fig. 1). Here, we report that this tripartite system enabled one to visualize the formation of ternary protein complexes in living cells with high spatial and temporal resolution.

**Figure 1.**
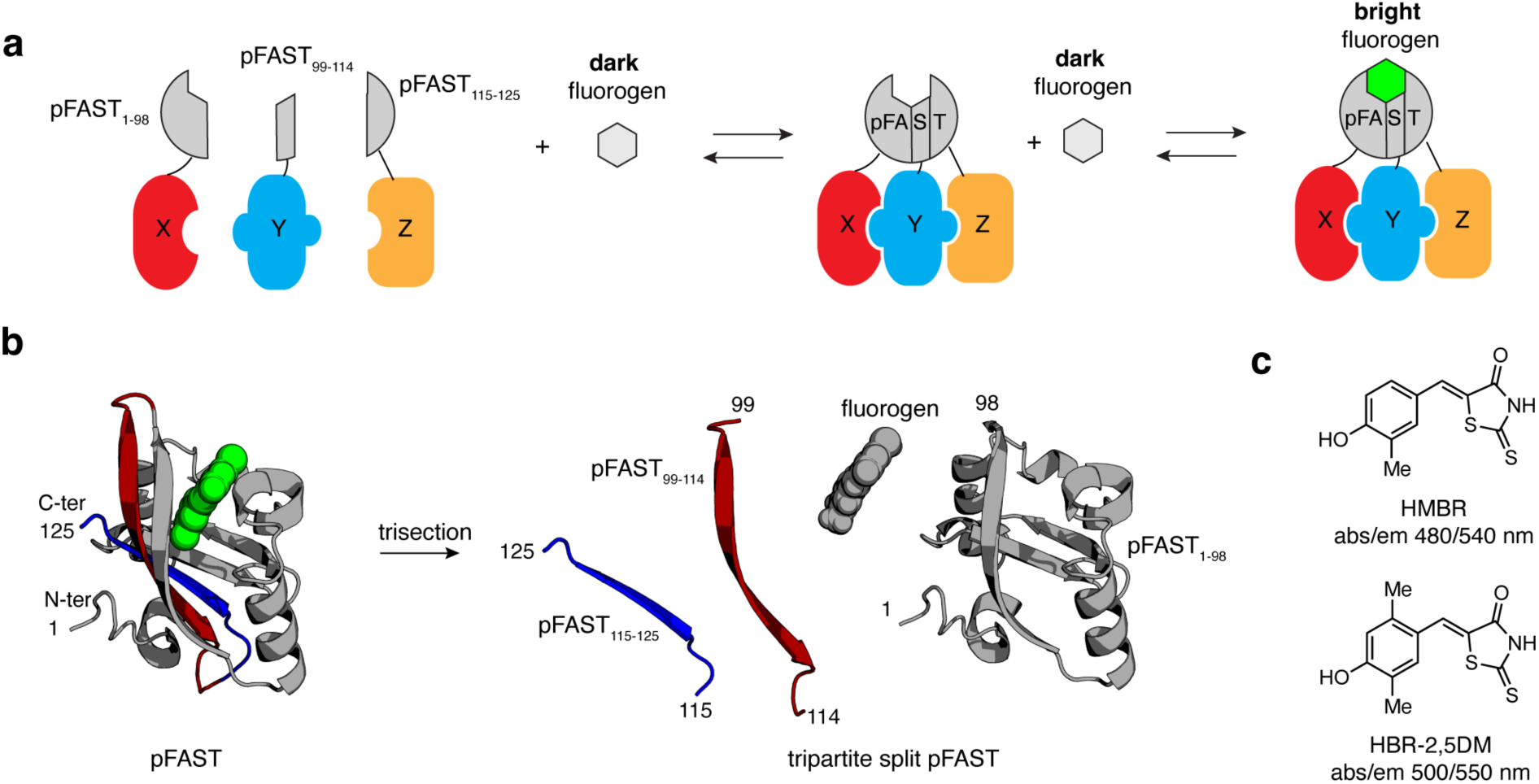
A complementation-based assay to visualize ternary protein interactions. **a** The interaction of three proteins is visualized through proximity-induced complementation of three complementary fragments of the chemogenetic fluorescent reporter pFAST. **b** Design of tripartite-split-pFAST through protein splitting at sites 98-99 and 114-115. The model of pFAST was generated in ref. ^27^. **c** Structures of the fluorogens 4-hydroxy-3-methylbenzylidene rhodanine (HMBR) and 4-hydroxy-2,5-dimethylbenzylidene rhodanine (HBR-2,5DM) used in this study.

## RESULTS

### New functional site for bisection of FAST

Bipartite-split-FAST was originally created through bisection of FAST between residues 114 and 115^8^, a split site originally used for the design of functional circular permutations of FAST variants^22^. In order to identify alternative split sites, we screened for alternative circular permutations of FAST. As the systematic and comprehensive study of all possible circular permutations of a protein, albeit feasible, can be tedious and time-consuming, we used Circular Permutation site predictor (CPred), a computational tool for predicting viable circular permutations in proteins^23^. In absence of crystal structure for FAST and its variants, we used the crystal structure of *Halorhodospira halophila* Photoactive Yellow Protein (PYP), the parental protein from which prototypical FAST was evolved^12^. We assumed that, because of the high sequence homology between PYP and FAST and their common structural features, the results from PYP analysis could be directly transposed to FAST. CPred provided a probability score for each position, allowing to identify several potential viable cleavage sites (Supplementary Fig. 1). After rational evaluation of the suggested regions, we retained six potential split sites with high probability scores: 24-25, 35-36, 59-60, 72-73, 89-90, 98-99 and 102-103, and generated the corresponding circular permutations by insertion of a ENLYFQGGSGG linker between FAST native termini (Fig. 2a). This linker constituted a relatively long and flexible linker, enabling to evaluate the viability of the circular permutation. The DNA sequences encoding the seven circular permutations were generated by polymerase chain reaction (PCR) from the sequence encoding a FAST tandem connected with a ENLYFQGGSGG linker (Fig. 2a). The circular permutations cpFAST(35-36) and cpFAST(102-103) could not be expressed and purified in *E. coli* in preliminary tests, and were thus not further studied. Among the five remaining cpFAST variants, only cpFAST(24-25) and cpFAST(98-99) were able to bind the fluorogenic chromophore 4-hydroxy-3-methylbenzylidene rhodanine (HMBR) and activate its fluorescence (Fig. 2a and Supplementary Figs. 2 & 4), however with lower fluorogen binding affinity than prototypical FAST.

**Figure 2.**
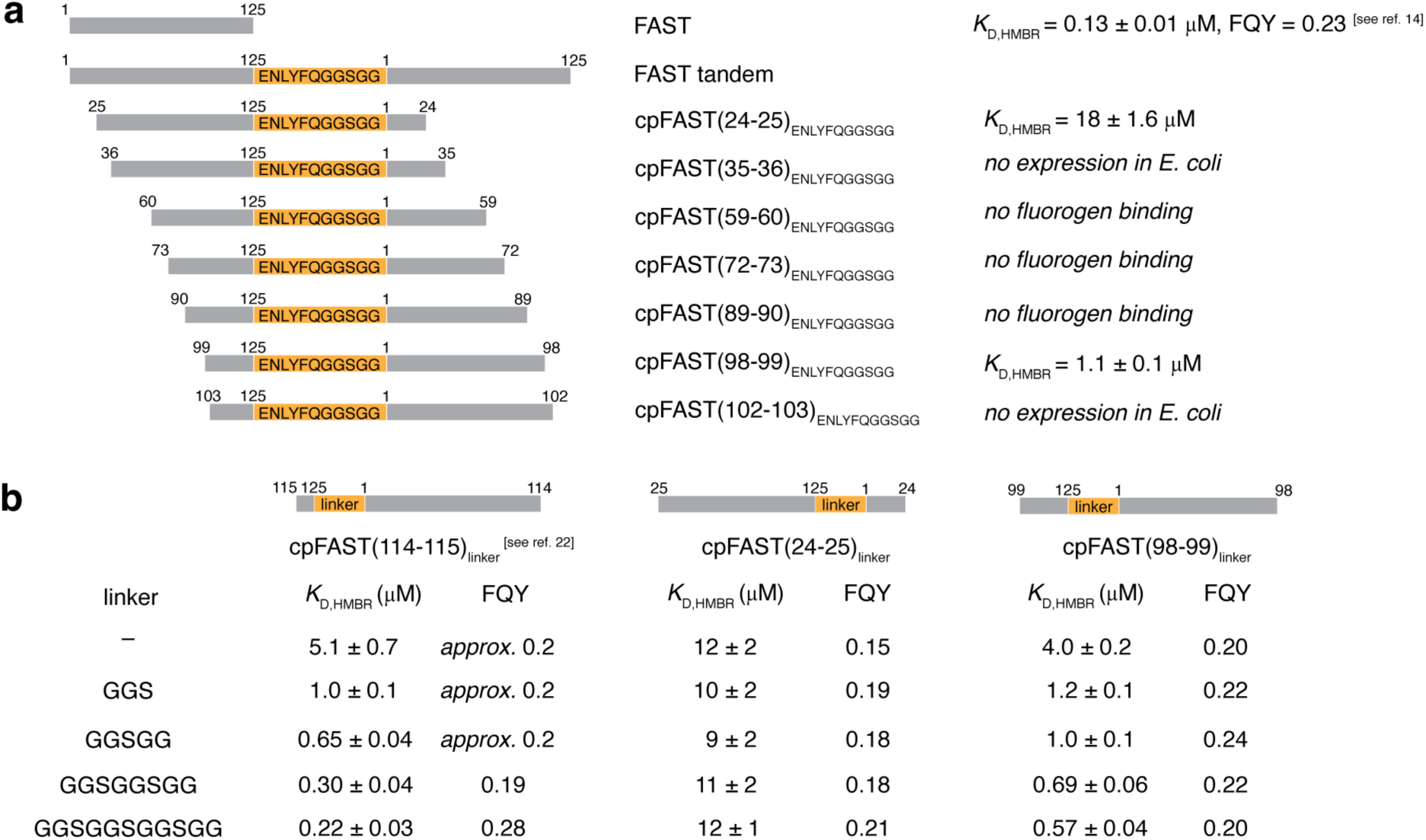
Circular permutations of FAST. **a** Sequences and properties of the different circular permutations of FAST screened during this study (see also Supplementary Fig. 2 for details). **b** Influence of the linker on the properties of cpFAST(24-25) and cpFAST(98-99). Are reported the thermodynamic dissociation constants and the fluorescence quantum yields (FQY) of the different cpFAST::HMBR assemblies (see also Supplementary Figs. 3 & 4). Previously reported data for cpFAST(114-115) are shown for comparison.

As the length of the linker connecting the native termini can play a role in the folding and activity, we next replaced the ENLYFQGGSGG linker by flexible GGS linkers of varying size from 11 to 3 residues. A version with no linker between the native termini was also tested for both circular permutations. All new variants were able to form a fluorescent assembly with HMBR and give good fluorescence quantum yield (Fig. 2b and Supplementary Figs. 3 & 4), however the effect of the linker length on the fluorogen binding affinity was notably different between the two circular permutations. In the case of cpFAST(98-99), increasing the linker length from 0 to 11 amino acids increased the fluorogen binding affinity, as shown by the dissociation constant *K*_D_ varying from 4.0 ± 0.2 to 0.6 ± 0.1 μM, reaching binding affinity close to that of FAST. These results suggested that the use of an 11 amino acid linker allows cpFAST(98-99) to adopt a conformation close to FAST, as previously observed for cpFAST(114-115)^20^. On the other hand, in the case of cpFAST(24-25), no change of affinity was observed when varying the linker length: *K*_D_ remained around 10 μM no matter the linker used. This binding affinity was close to that observed for nanoFAST (a.k.a. FAST_27-125_) (Supplementary Fig. 5), a version of FAST in which the first 26 residues are truncated^24^. This suggested that only the domain FAST_25-125_ was properly folded and functional within the different cpFAST(24-25) or that the two domains FAST_25-125_ and FAST_1-24_ were folding independently without interacting together. In both cases, these results precluded to use the split site 24-25 for the generation of a functional split version. We thus focused our study only on the bisection of FAST at position 98-99.

To evaluate whether the two fragments FAST_1-98_ and FAST_99-125_ could complement and form a functional system when in close proximity, the two fragments were fused respectively to the N-terminus of the FRB (FKBP-rapamycin binding) domain of mTOR (mechanistic target of rapamycin), and to the C-terminus of FKBP (FK506-binding protein)^25,26^ (Fig. 3b,c). As FRB and FKBP can interact together in presence of rapamycin, this pair of proteins enables to evaluate both interaction-dependent complementation in the presence of rapamycin and self-complementation in its absence^8^. The interaction-induced pre-organization of the two FAST’s fragments can increase the efficiency of their complementation with the fluorogen by reducing this three-body system to a ‘virtual’ two-body system (Fig. 3a). In order to evaluate the complementation efficiency in the two scenarios, FAST_1-98_–FRB and FKBP–FAST_99-125_ were co-expressed in HEK293T cells using two bicistronic plasmids vectors that allowed also the expression of mTurquoise2 and iRFP670, respectively, used here as transfection reporters (Supplementary Fig. 6). As the complementation efficiency of split-FAST(98-99) depends on the fluorogen concentration (Fig. 3a)^11^, we quantified it by flow cytometry in the presence or the absence of rapamycin at various concentrations of the fluorogen HMBR. This experiment enabled us to determine the effective concentration of fluorogen for half-maximum complementation in the absence (EC_50,–interaction_) or the presence (EC_50,+interaction_) of rapamycin (Fig 3c). These values can thus be used to evaluate the efficacy of the self-complementation and of the interaction-dependent complementation: the higher the EC_50,–interaction_, the lower the self-complementation, and the lower the EC_50,+interaction_, the higher the interaction-dependent complementation efficacy. Note that the obtained complementation curves enable also to identify the optimal fluorogen concentration to maximize the interaction-induced complementation while minimizing self-complementation. We found an EC_50,– interaction_ = 31 ± 4 μM and an EC_50,+interaction_ = 7.5 ± 0.7 μM, demonstrating that the complementation of split-FAST(98-99) is favored when FRB and FKBP interact together, even though the value of EC_50,–interaction_ revealed that the two fragments could complement to a certain extent in the absence of interaction at high concentration of fluorogen. Globally, the behavior of split-FAST(98-99) was very similar to the one observed for regular split-FAST(114-115) (made of FAST_1-114_ and FAST_115-125_) fragments^8,11^, suggesting that splitting FAST at position 98-99 led to a functional split system.

**Figure 3.**
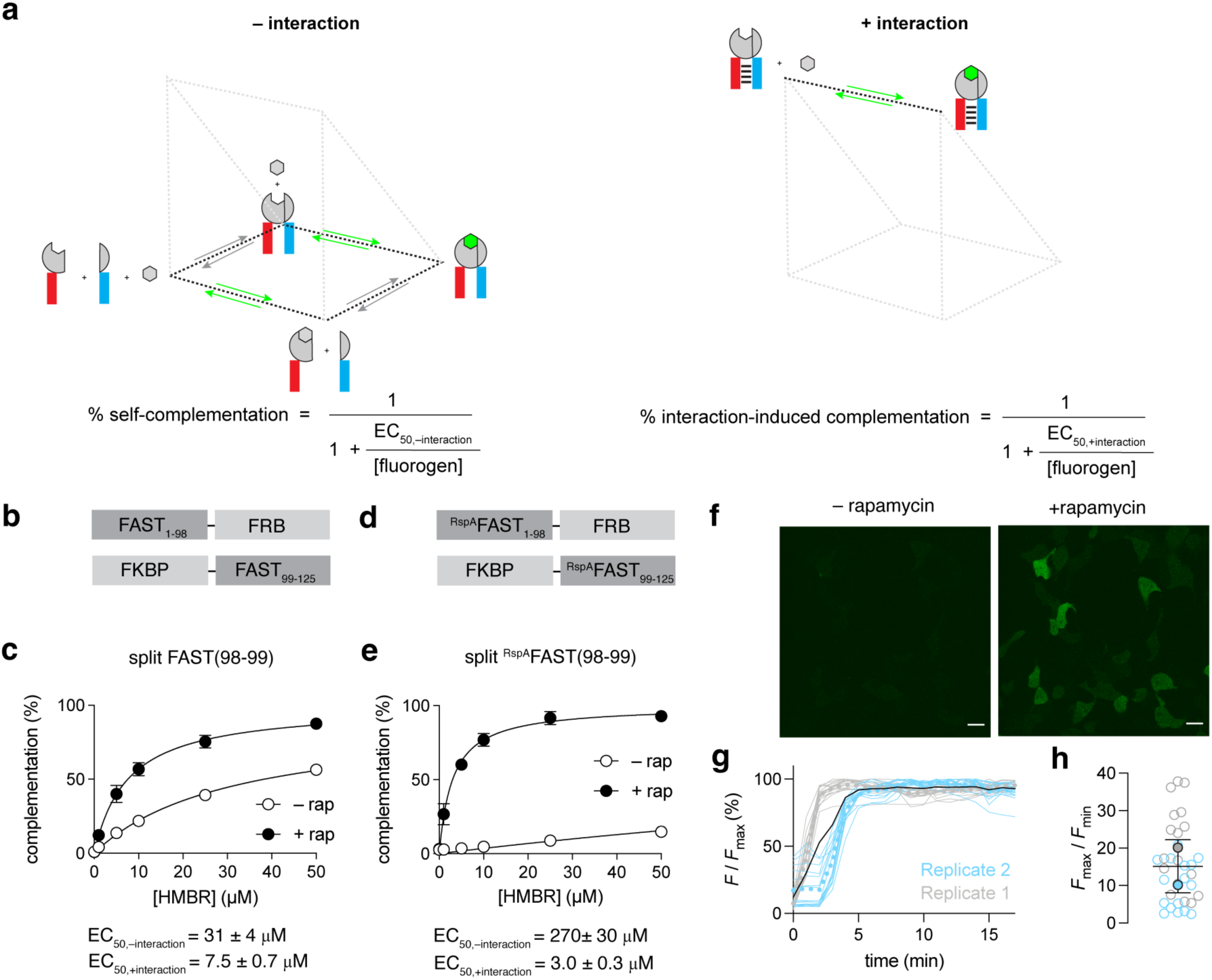
Evaluation of new split site 98-99. **a** Model for the self-complementation and interaction-dependent complementation of bipartite split-FAST. Evaluation of the complementation efficiency in function of the fluorogen concentration enables to extract the effective fluorogen concentration for half maximal complementation in the absence of interaction (EC_50,–interaction_) and in the presence of interaction (EC_50_,_+interaction_). The EC_50,–interaction_ and EC_50,+interaction_ values can be used to evaluate the efficacy of the self-complementation and of the interaction-dependent complementation: the higher the EC_50,–interaction_, the lower the self-complementation, and the lower the EC_50,+interaction_, the higher the interaction-dependent complementation efficacy. The change in EC_50_ allows us furthermore to evaluate the dynamic range of the system, the higher the change, the better the dynamic range, and thus the greater the system for the monitoring of interactions. **b-e** Normalized average fluorescence of about 20,000 HEK293T cells co-expressing the FK506-binding protein (FKBP) fused to the N-terminus of FAST_99-125_ (**b,c**) or ^RspA^FAST_99-125_ (**d,e**) and the FKBP-rapamycin-binding domain of mammalian target of rapamycin (FRB) fused to the C-terminus of FAST_1-98_ (**b,c**) or ^RspA^FAST_1-98_ (**d,e**) treated without or with 500 nM of rapamycin, and with 0, 1, 5, 10, 25 or 50 μM of HMBR. Data in **c,e** represent the mean ± SD of three independent experiments. The concentrations of fluorogen for half maximal complementation in the absence (EC_50,–interaction_) or the presence (EC_50,+interaction_) of rapamycin are given. **f-h** HEK293T cells expressing ^RspA^FAST_1-98_-FRB and FKBP-^RspA^FAST_99-125_ were treated with 10 μM HMBR. Cells were imaged by time-lapse confocal microscopy after addition of 500 nM of rapamycin. Experiments were repeated two times with similar results. **f** Representative micrographs before and after addition of rapamycin (see also Supplementary Movie 1). Scale bars 20 μm. **g** Temporal evolution of the fluorescence signal (*F*/*F*_max_) after addition of rapamycin n = 33 cells from two independent experiments. *F* is the fluorescence intensity at time t, and *F*_max_ is the maximal fluorescence intensity reached during the time-lapse. **h** Fluorescence fold increase (*F*_max_/*F*_min_) upon addition of rapamycin. *F*_min_ is the fluorescence intensity at time t = 0, and *F*_max_ is the maximal fluorescence intensity reached during the time-lapse. **g,h** Each cell is color-coded according to the biological replicate it came from. **g** The dot lines represent the mean value of each biological replicate, while the black line represents the mean of the two biological replicates. **h** The solid circles correspond to the mean of each biological replicate. The black line represents the mean ± SD of the two biological replicates.

We recently generated split-FAST2, a version of split-FAST(114-115) with reduced self-complementation and improved interaction-dependent complementation. This improved system was generated by splitting ^RspA^FAST, a FAST variant generated from a PYP ortholog of *Rheinheimera sp.* A13L (RspA), displaying the same length as FAST and 78% sequence identity (See Supplementary Fig. 9 for the sequence of FAST and RspA-FAST, and a summary of the properties of their full-length and split versions). split-FAST2 was obtained by splitting ^RspA^FAST between the positions 114 and 115, used to generate split-FAST, giving two fragments ^RspA^FAST_1-114_ and ^RspA^FAST_115-125_^11^. We applied this orthology-based engineering strategy to split-FAST(98-99). We split ^RspA^FAST at position 98-99 to generate two fragments, ^RspA^FAST_1-98_ and ^RspA^FAST_99-125_, and characterized their complementation fusing them to FRB and FKBP (Fig. 3d,e). Flow cytometry analysis of the complementation of ^RspA^FAST_1-98_–FRB and FKBP–^RspA^FAST_99-125_ at various HMBR concentrations, and in the absence or the presence of rapamycin showed that the use of ^RspA^FAST fragments enabled to increase the efficacy of interaction-dependent complementation by 2.5-fold (EC_50,+interaction_ = 3.0 ± 0.3 μM), and dramatically reduced self-complementation (EC_50,– interaction_ = 270 ± 30 μM), leading to a significant increase of dynamic range (100-fold). Monitoring the rapamycin-induced complementation of ^RspA^FAST_1-98_–FRB and FKBP– ^RspA^FAST_99-125_ in HEK293T cells treated with 10 μM HMBR by confocal microscopy confirmed that ^RspA^FAST_1-98_ and ^RspA^FAST_99-125_ show minimal self-complementation in the presence of HMBR and efficiently complement in a proximity-dependent manner (Fig. 3f-h and Supplementary Movie 1). Overall, this set of experiments showed that the site 98-99 could be a functional bisection site for the generation of a bipartite-split-FAST.

### Trisection of FAST variants

With the final goal of generating a tripartite-split-FAST able to report on the proximity of three proteins, we combined the sites 98-99 and 114-115 to split ^RspA^FAST into three fragments: ^RspA^FAST_1-98_, ^RspA^FAST_99-114_ and ^RspA^FAST_115-125_. To evaluate the potential of these three fragments to complement into a functional reporter capable of binding fluorogens when in close proximity, we first fused two of the three fragments to the N- and C-termini of FRB and the third fragment to FKBP. We took advantage of the structure of the FRB-rapamycin-FKBP complex to identify the right topology to use. After predicting tridimensional structures with Alphafold^27^ (Supplementary Fig. 7), we decided to fuse ^RspA^FAST_99-114_ and ^RspA^FAST_1-98_ to the N- and C-termini of FRB, respectively, and ^RspA^FAST_115-125_ to the C-terminus of FKBP (Fig. 4a). We evaluated their complementation in HEK293T cells by flow cytometry in the absence or presence of rapamycin at various concentrations of HMBR (Fig. 4b). In the absence of rapamycin, no complementation was observed, no matter the concentration of HMBR used. In the presence of rapamycin, complementation could be observed at high HMBR concentrations, but with relatively modest efficiency compared to the bipartite versions. The modest complementation efficiency could result from an improper reassembly or imperfect folding of the three fragments resulting in a loss of fluorogen binding ability/affinity, limiting thus the formation of the quaternary fluorescent assembly. When using 4-hydroxy-2,5-dimethylbenzylidene rhodanine (HBR-2,5DM), a fluorogen that binds FAST tighter than HMBR^14^, we observed a significant improvement of the interaction-dependent complementation efficacy, reaching an EC_50,+ interaction_ = 8.1 ± 0.2 μM, while keeping minimal self-complementation (Fig. 4c). This result supported the idea that the use of highly affine fluorogens favors the formation of the quaternary fluorescent assembly by compensating the loss of entropy resulting from the trisection. We hypothesized that efficient complementation required additional binding energy to counteract the unfavorable entropy change associated with assembling together the three fragments and the fluorogen into a quaternary fluorescent assembly. If so, additional binding energy could favor the interactions of the three fragments or the fluorogen binding or both.

**Figure 4.**
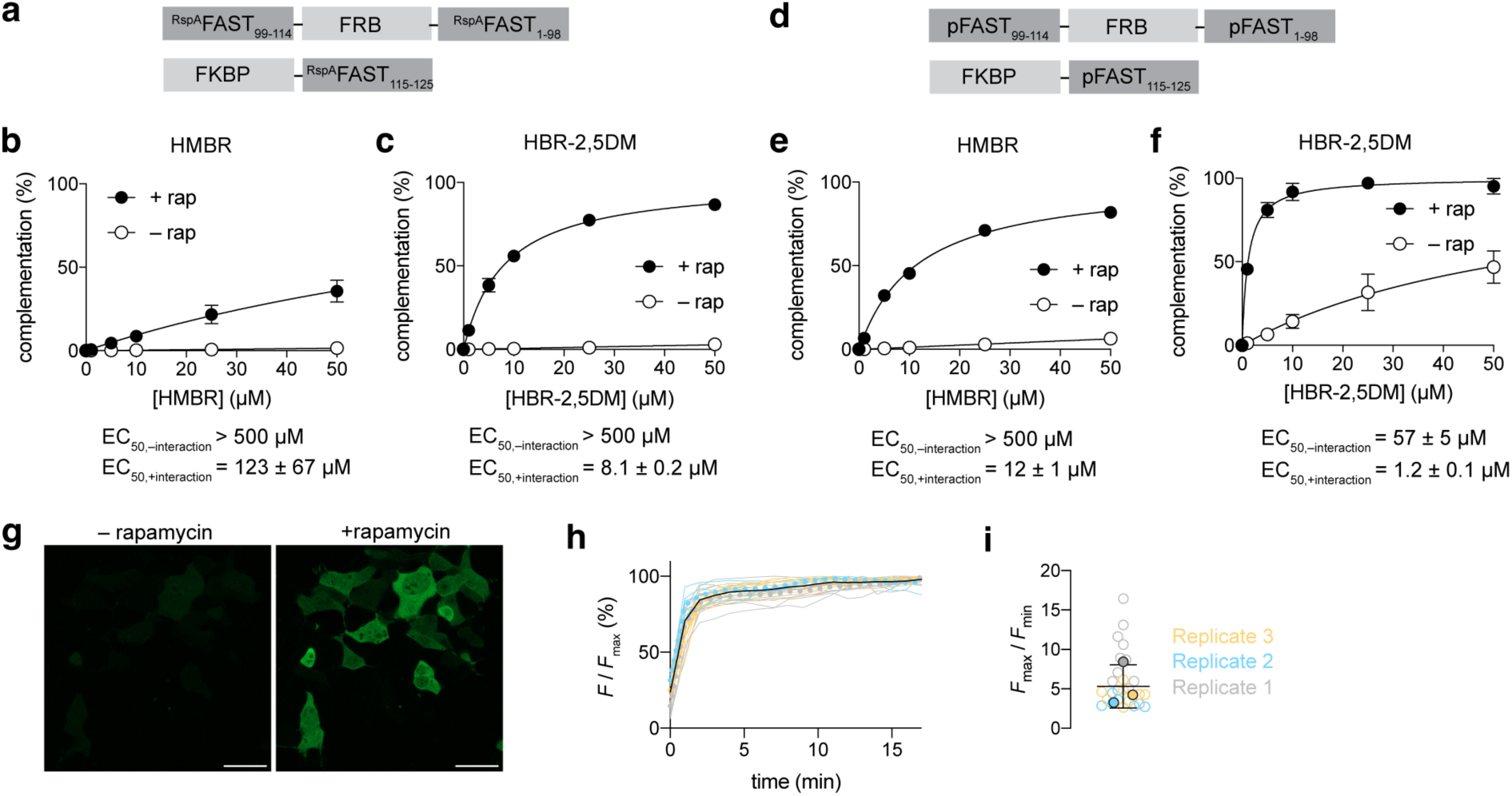
Evaluation of tripartite split systems. **a-f** Normalized average fluorescence of about 20,000 HEK293T cells co-expressing the pairs ^RspA^FAST_99-114_–FRB–^RspA^FAST_1-98_ / FKBP–^RspA^FAST_115-125_ (**a-c**) or pFAST_99-114_–FRB–pFAST_1-98_ / FKBP–pFAST_115-125_ (**d-f**) treated without or with 500 nM of rapamycin, and with 0, 1, 5, 10, 25 or 50 μM of HMBR (**b,e**) or HBR–2,5DM (**c,f**). Data represent the mean ± SD of three independent experiments (**b,f**) and two independent experiments (**c,e**). The concentrations of fluorogen for half maximal complementation in the absence (EC_50,–interaction_) or the presence (EC_50,+interaction_) of rapamycin are given. **g-i** HEK293T cells expressing pFAST_99-114_–FRB–pFAST_1-98_ and FKBP–pFAST_115-125_ were treated with 5 μM HBR-2,5DM. Cells were imaged by time-lapse confocal microscopy after addition of 500 nM of rapamycin. Experiments were repeated three times with similar results. **g** Representative micrographs before and after addition of rapamycin (see also Supplementary Movie 2). Scale bars 20 μm. **h** Temporal evolution of the fluorescence signal (*F*/*F*_max_) after addition of rapamycin n = 26 cells from three independent experiments. *F* is the fluorescence intensity at time t, and *F*_max_ is the maximal fluorescence intensity reached during the time-lapse. **i** Fluorescence fold increase (*F*_max_/*F*_min_) upon addition of rapamycin. *F*_min_ is the fluorescence intensity at time t = 0, and *F*_max_ is the maximal fluorescence intensity reached during the time-lapse. **h,i** Each cell is color-coded according to the biological replicate it came from. **h** The dot lines represent the mean value of each biological replicate, while the black line represents the mean of the three biological replicates. **i** The solid circles correspond to the mean of each biological replicate. The black line represents the mean ± SD of the three biological replicates.

We recently described a bipartite split system with high self-complementation, i.e. capable of complementation in the presence of fluorogen even in the absence of interaction. This system allowed us to develop a fluorogenic chemically induced dimerization tool to induce and visualize the proximity of two proteins^28^. This CATCHFIRE (Chemically Assisted Tethering of CHimera by Fluorogenic Induced Recognition) approach was made through bisection of pFAST at position 114-115, generating two fragments pFAST_1-114_ (a.k.a. ^FIRE^mate) and pFAST_115-125_ (a.k.a. ^FIRE^tag) (See Supplementary Fig. 9 for the sequence of pFAST and a summary of the properties of its full-length and split versions). pFAST is a FAST variant able to bind fluorogenic chromophores with spectral properties covering the entire visible spectrum^29^. pFAST was shown to display higher conformational rigidity, resulting overall in higher fluorogen binding affinity. We assume that higher folding and binding energy provides the driving force for the self-complementation of the fluorescent ternary assembly in the case of CATCHFIRE. We thus hypothesized that trisection of pFAST could result in a tripartite system with higher complementation efficiency. To test this idea, we fused pFAST_99-114_ and pFAST_1-98_ to the N- and C-termini of FRB, respectively, and pFAST_115-125_ to the C-terminus of FKBP (Supplementary Fig. 8), and evaluated their complementation in HEK293T cells by flow cytometry as previously done for the tripartite-split-^RspA^FAST (Fig. 4d-f). The analysis was done in the absence or the presence of rapamycin at various concentrations of HMBR or HBR-2,5DM. With HMBR, almost no self-complementation could be detected in the absence of rapamycin, however we observed an efficient complementation when inducing the FRB-FKBP interaction by addition of rapamycin, reaching an EC_50,+ interaction_ = 12 ± 1 μM. With HBR-2,5DM, the efficacy of interaction-dependent complementation was 10-fold higher (EC_50,+ interaction_ = 1.2 ± 0.1 μM), supporting the idea that the use of fluorogens with high binding affinity favors the formation of the quaternary fluorescent assembly. Although the system could partially self-complement at high HBR-2,5DM concentrations, self-complementation remains very low at low concentrations, allowing to detect the rapamycin-induced interaction of FRB and FKBP with high dynamic range. For instance, at [HBR-2,5DM] = 5 μM, complementation reaches 81% with rapamycin and less than 6.5% without, resulting in a 12-fold fluorescence increase upon formation of the interaction. Monitoring the rapamycin-induced complementation of pFAST_99-114_–FRB–pFAST_1-98_ and FKBP–pFAST_114-125_ in HEK293T cells treated with 5 μM HBR-2,5DM by confocal microscopy confirmed that the tripartite-split-pFAST showed minimal self-complementation in presence of HBR-2,5DM and could efficiently complement in a proximity-dependent manner (Fig. 4g-i and Supplementary Movie 2). Kinetic analysis showed that the rapamycin-induced formation of the quaternary green fluorescent assembly was rapid and occurred with half-time < 2 min. This timescale was comparable to that observed with bipartite-split-FASTs^8,11^.

The experiments described above allowed us to demonstrate that the three fragments resulting from the splitting of pFAST at positions 98-99 and 114-115 were able to efficiently complement with HMBR or HBR-2,5DM when brought into close proximity. However, we used a model system in which the fragments pFAST_1-98_ and pFAST_99-114_ were fused to the same protein, which already preorganized them to interact. To demonstrate that the tripartite split system could selectively complement when the three fragments are fused to three different proteins that interact together (Fig. 5a), we used the chemically inducible heterotrimerization (CIT) system recently developed from the FKBP-FRB-rapamycin system^30^. In this CIT system, trimerization of a bipartite-split-FRB with full-length FKBP can be induced by addition of rapamycin. We fused pFAST_99-114_ at the N-terminus of the N-terminal fragment of split-FRB (pFAST_99-114_–^N^FRB) and pFAST_1-98_ at the C-terminus of its C-terminal fragment (^C^FRB–pFAST_1-98_), and kept pFAST_115-125_ fused to the C-terminus of FKBP (Fig. 5b). To express pFAST_99-114_–^N^FRB and ^C^FRB–pFAST_1-98_ with a single plasmid, we constructed a bicistronic vector containing a viral P2A cleavage sequence in between the sequences coding for the two proteins. We co-expressed the three proteins pFAST_99-114_–^N^FRB, ^C^FRB–pFAST_1-98_ and FKBP–pFAST_115-125_ in HEK293T cells, treated cells with HBR-2,5DM, and evaluated the complementation efficiency of the tripartite split system in the absence or presence of rapamycin by flow cytometry (Fig. 5c). Various concentrations of HBR-2,5DM were tested to evaluate the role of fluorogen concentration on complementation efficiency. As for the bipartite split-FAST, the complementation efficiency of the tripartite split-FAST depends on the fluorogen concentration (Fig. 5a). Similarly, the interaction-induced pre-organization of the three FAST’s fragments reduces the four-body system to a two-body system, and increases consequently complementation efficacy (Fig. 5a). We used the concentration of fluorogen for half maximal complementation as a proxy of the efficiency of formation of the quaternary fluorescent assembly (Fig. 5a). We observed that complementation efficiency increased by almost two orders of magnitude when the three fragments are forced into close proximity by chemically induced trimerization. Our analysis showed that it is possible to find fluorogen concentrations for which interaction-dependent complementation is maximal or quasi-maximal, while self-complementation remains minimal (i.e. below 10%), ensuring the detection of ternary interaction with high dynamic range.

**Figure 5.**
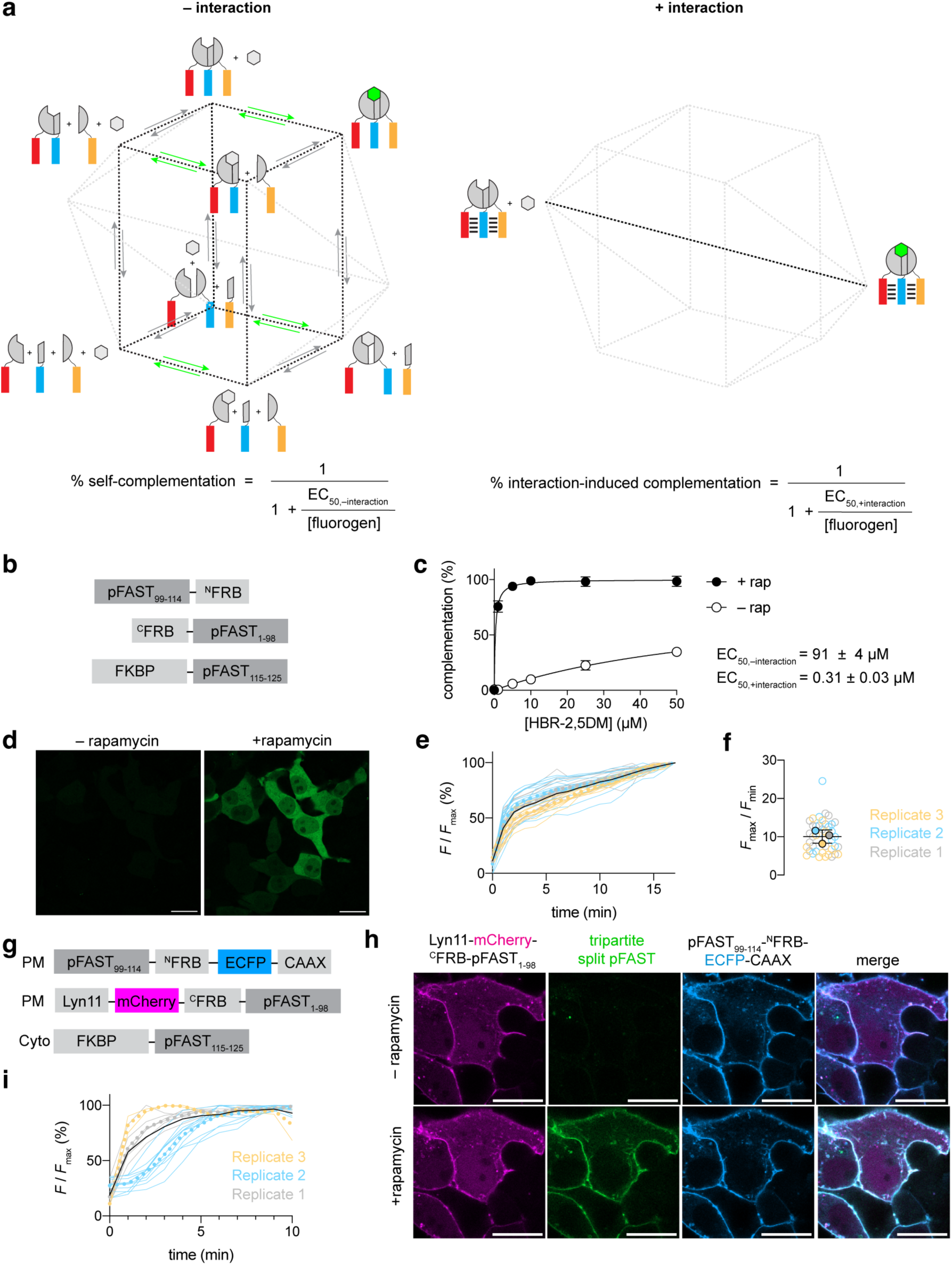
Detection of ternary protein interactions. **a** Model for the self-complementation and interaction-dependent complementation of tripartite split-FAST. Evaluation of the complementation efficiency in function of the fluorogen concentration enables to extract the effective fluorogen concentration for half maximal complementation in the absence of interaction (EC_50_,_–interaction_) and in the presence of interaction (EC_50_,_+interaction_). The EC_50,–interaction_ and EC_50,+interaction_ values can be used to evaluate the efficacy of the self-complementation and of the interaction-dependent complementation: the higher the EC_50,–interaction_, the lower the self-complementation, and the lower the EC_50,+interaction_, the higher the interaction-dependent complementation efficacy. The change in EC_50_ allows us furthermore to evaluate the dynamic range of the system, the higher the change, the better the dynamic range, and thus the greater the system for the monitoring of interactions. **b,c** Normalized average fluorescence of about 20,000 HEK293T cells co-expressing the triplet pFAST_99-114_–^N^FRB, ^C^FRB–pFAST_1-98_ and FKBP–pFAST_115-125_ treated without or with 500 nM of rapamycin, and with 0, 1, 5, 10, 25 or 50 μM of HBR– 2,5DM. Data represent the mean ± SD of two independent experiments. The concentrations of fluorogen for half maximal complementation in absence (EC_50,–interaction_) or presence (EC_50,+interaction_) of rapamycin are given. **d-f** HEK293T cells expressing the triplet pFAST_99-114_–^N^FRB, ^C^FRB–pFAST_1-98_ and FKBP– pFAST_115-125_ were treated with 10 μM HBR-2,5DM. Cells were imaged by time-lapse confocal microscopy after addition of 500 nM of rapamycin. Experiments were repeated three times with similar results. **d** Representative micrographs before and after addition of rapamycin (see also Supplementary Movie 3). Scale bars 20 μm. **e** Temporal evolution of the fluorescence signal (*F*/*F*_max_) after addition of rapamycin n = 45 cells from three independent experiments. *F* is the fluorescence intensity at time t, and *F*_max_ is the maximal fluorescence intensity reached during the time-lapse. **f** Fluorescence fold increase (*F*_max_/*F*_min_) upon addition of rapamycin. *F*_min_ is the fluorescence intensity at time t = 0, and *F*_max_ is the maximal fluorescence intensity reached during the time-lapse. **g-i** HEK293T cells expressing the triplet pFAST_99-114_–^N^FRB–ECFP–CAAX, Lyn11-mCherry–^C^FRB–pFAST_1-98_ and FKBP–pFAST_115-125_ were treated with 10 μM HBR-2,5DM. Cells were imaged by time-lapse confocal microscopy after addition of 500 nM of rapamycin. Experiments were repeated three times with similar results. **h** Representative micrographs before and after addition of rapamycin (see also Supplementary Movie 4). Scale bars 20 μm. **i** Temporal evolution of the fluorescence signal (*F*/*F*_max_) after addition of rapamycin n = 20 cells from three independent experiments. *F* is the fluorescence intensity at time t, and *F*_max_ is the maximal fluorescence intensity reached during the time-lapse. **e,f,i** Each cell is color-coded according to the biological replicate it came from. **e,i** The dot lines represent the mean value of each biological replicate, while the black line represents the mean of the three biological replicates. **f** The solid circles correspond to the mean of each biological replicate. The black line represents the mean ± SD of the three biological replicates.

Next, we studied the kinetics and dynamics of complementation in living cells by time-lapse fluorescence confocal microscopy (Fig. 5d-f and Supplementary Movie 3). We monitored the formation of the quaternary fluorescent assembly upon rapamycin-induced trimerization of pFAST_99-114_-^N^FRB, ^C^FRB-pFAST_1-98_ and FKBP-pFAST_115-125_ in HEK293T cells pretreated with 10 μM HBR-2,5DM. In the absence of rapamycin, almost no fluorescence was detectable in agreement with the low self-complementation of the tripartite-split-pFAST at this fluorogen concentration. Addition of rapamycin led to the rapid formation of the quaternary fluorescent assembly, demonstrating the ability of the tripartite-split-pFAST to visualize the formation of a ternary protein complex. In average, a tenfold fluorescence increase was observed upon rapamycin-induced trimerization of ^N^FRB, ^C^FRB and FKBP.

### Visualization of ternary protein interaction at the plasma membrane

As many ternary protein interactions occur at the plasma membrane (PM), we next asked if tripartite split-pFAST could detect the interaction between two proteins anchored at the PM inner face and a third protein localized in the cytosol. We fused ^C^FRB-pFAST_1-98_ to a Lyn11 membrane-anchoring peptide sequence from Lyn kinase and the red fluorescent protein mCherry (Lyn11–mCherry– ^C^FRB-pFAST_1-98_), and pFAST_99-114_-^N^FRB to a CAAX sequence and the enhanced cyan fluorescent protein (ECFP) (pFAST_99-114_-^N^FRB–ECFP–CAAX) (Fig. 5g-i). The fluorescent proteins mCherry and ECFP allowed to verify the proper localization of the proteins at the PM. We expressed in HEK293T cells the two proteins together with FKBP-pFAST_115-125_ as a cytosolic partner and treated cells with 10 μM HBR-2,5DM. The addition of rapamycin led to the rapid appearance of green fluorescence at the PM (Fig. 5h,i and Supplementary Movie 4), in agreement with the rapamycin-induced interaction of the three proteins, demonstrating the ability of tripartite-split-pFAST to sense ternary interactions at the PM.

### Visualization of ternary protein interactions at organelle membrane junctions

As the three fragments of tripartite-split-pFAST can complement into a functional reporter when in close proximity, we then asked if tripartite-split-pFAST could detect the recruitment of proteins at organelle membrane contact sites or junctions. In eukaryotic cells, organelles do not work as isolated structures, but form a dynamic, interconnected network which can be modulated according to cellular needs^2^. Close contacts between organellar membranes, so-called membrane contact sites (MCSs), are involved in key cellular functions such as intracellular signaling, autophagy, lipid metabolism, membrane dynamics, organelle trafficking and biogenesis^1^. The formation and dynamics of MCSs are regulated by protein-protein and protein-lipid interactions, modulated by specific molecular tethers and actuators localized at membrane contacts and junctions^2,31^. Molecular tools enabling to visualize the proximity of three proteins can thus enable to study the recruitment of cytosolic molecular effectors at organelle membrane contact sites and junctions with high spatial and temporal resolution. As a proof of principle, we used the tripartite-split-pFAST to visualize the recruitment of a cytosolic protein at mitochondria (MT)–endoplasmic reticulum (ER), ER–PM and MT– PM membrane contact sites/junctions chemically induced using the split-FRB-rapamycin-FKBP CIT tool^30^. MT-ER, ER-PM and MT-PM membrane contact sites/junctions were chosen as archetypes of membrane contact sites because of their biological relevance in the regulation of various biological processes^1^.

To visualize the recruitment of a cytosolic protein at the MT–PM membrane contact sites/junctions, we co-expressed FKBP–pFAST_115-125_ together with MT-targeted mCherry–^C^FRB–pFAST_1-98_ (through fusion to the mitochondrial import receptor TOM20, giving TOM20–mCherry–^C^FRB–pFAST_1-98_), and PM-targeted pFAST_99-114_–^N^FRB–ECFP (through fusion to a CAAX sequence, giving pFAST_99-114_– ^N^FRB–ECFP–CAAX) in HeLa cells (Fig. 6a-c). Cells were treated with HBR-2,5DM before addition of rapamycin. After addition of rapamycin, we observed a significant increase of the image correlation coefficient between the MT and PM signals (Fig. 6c), in agreement with the formation of MT-PM junctions. We also observed the appearance of a green fluorescent signal that colocalized with patches formed between MT and PM (Fig. 6b and Supplementary Movie 5), suggesting that tripartite split-pFAST can enable to visualize the recruitment of a cytosolic protein at the MT-PM junctions.

**Figure 6.**
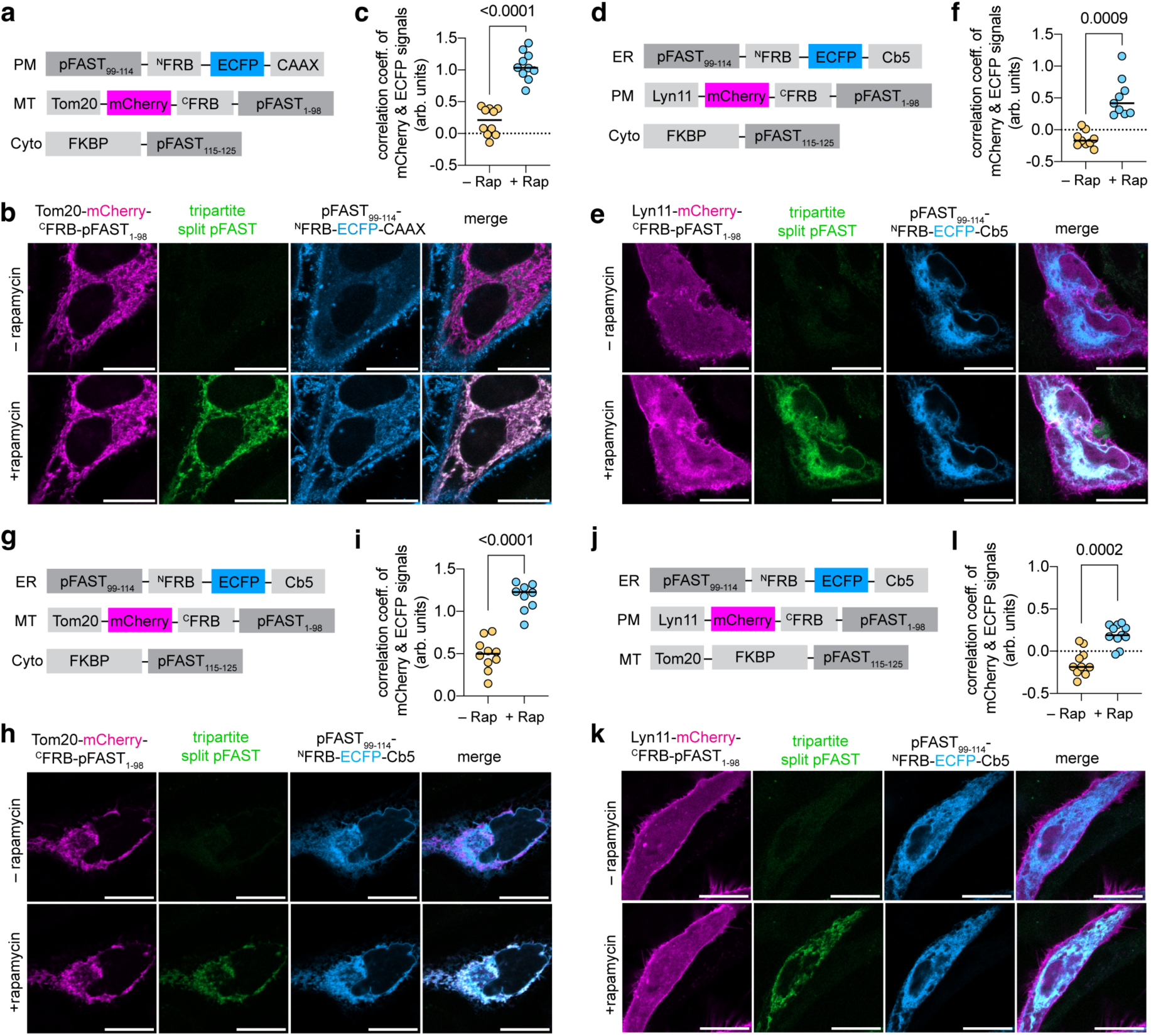
Visualization of ternary protein interactions at the junctions of organelles. **a-c** HeLa cells co-expressing pFAST_99-114_–^N^FRB–ECFP–CAAX, Tom20-mCherry–^C^FRB–pFAST_1-98_ and FKBP– pFAST_115-125_ were treated with 10 μM HBR-2,5DM. Cells were imaged by time-lapse confocal microscopy after addition of 500 nM of rapamycin. Experiments were repeated three times with similar results. **b** Representative micrographs before and after addition of rapamycin (see also Supplementary Movie 5). **c** Correlation of the ECFP and mCherry signals before and after rapamycin addition. **d-f** HeLa cells co-expressing pFAST_99-114_–^N^FRB–ECFP–Cb5, Lyn11-mCherry–^C^FRB–pFAST_1-98_ and FKBP– pFAST_115-125_ were treated with 10 μM HBR-2,5DM. Cells were imaged by time-lapse confocal microscopy after addition of 500 nM of rapamycin. Experiments were repeated three times with similar results. **e** Representative micrographs before and after addition of rapamycin (see also Supplementary Movie 6). **f** Correlation of the ECFP and mCherry signals before and after rapamycin addition. **c** Correlation of the EGFP and mCherry signals before and after rapamycin addition. **g-i** HeLa cells co-expressing pFAST_99-114_–^N^FRB–ECFP–Cb5, Tom20-mCherry–^C^FRB–pFAST_1-98_ and FKBP–pFAST_115-125_ were treated with 10 μM HBR-2,5DM. Cells were imaged by time-lapse confocal microscopy after addition of 500 nM of rapamycin. Experiments were repeated three times with similar results. **h** Representative micrographs before and after addition of rapamycin (see also Supplementary Movie 7). **i** Correlation of the EGFP and mCherry signals before and after rapamycin addition. **j-l** HeLa cells co-expressing pFAST_99-114_–^N^FRB–ECFP–Cb5, Lyn11-mCherry–^C^FRB–pFAST_1-98_, and Tom20–FKBP– pFAST_115-125_ were treated with 10 μM HBR-2,5DM. Cells were imaged by time-lapse confocal microscopy after addition of 500 nM of rapamycin. Experiments were repeated three times with similar results. **k** Representative micrographs before and after addition of rapamycin (see also Supplementary Movie 8). **l** Correlation of the EGFP and mCherry signals before and after rapamycin addition. **c,f,i,l** Data represent the mean of n = 10 (c), 9 (f), 10 (i), 11 (l) from three independent experiments. Two-tailed Student’s t-test assuming equal variance was used to compare correlations before and after addition rapamycin. Scale bars 20 μm.

We next applied tripartite split-pFAST to visualize the recruitment of a cytosolic protein at the ER–PM membrane contact sites/junctions (Fig. 6d-f). FKBP–pFAST_115-125_ was co-expressed together with PM-targeted mCherry–^C^FRB–pFAST_1-98_ (through fusion to a Lyn11 membrane-anchoring sequence, giving Lyn11–mCherry-^C^FRB– pFAST_1-98_) and ER-targeted pFAST_99-114_–^N^FRB–ECFP (through fusion to the cytochrome b5 (Cb5), giving pFAST_99-114_–^N^FRB–ECFP–Cb5) in HeLa cells. Addition of rapamycin to HBR-2,5-DM-pretreated cells led to a significant increase of the image correlation coefficient between the ER and PM signals (in agreement with the formation of ER-PM junctions) (Fig. 6f), and to the appearance of green fluorescent signal that colocalized with patches formed between PM and ER (Fig. 6e and Supplementary Movie 6), suggesting that tripartite-split-pFAST enabled to detect the recruitment of a cytosolic protein at the ER-PM junctions.

Similar conclusions were drawn when using tripartite-split-pFAST to monitor the recruitment of FKBP–pFAST_115-125_ at the ER–MT contact sites/junctions (Fig. 6g-i). Addition of rapamycin to HBR-2,5-DM-treated HeLa cells co-expressing FKBP– pFAST_114-115_ together with ER-targeted pFAST_99-114_–^N^FRB–ECFP–Cb5 and MT-targeted TOM20–mCherry–^C^FRB–pFAST_1-98_ led to a significant increase of the image correlation coefficient between the ER and MT signals (in agreement with the formation of ER-MT junctions) (Fig. 6i), and the appearance of green fluorescent signal that colocalized with patches between ER and MT (Fig. 6h and Supplementary Movie 7), in agreement with the assembly of tripartite-split-pFAST upon recruitment of the cytosolic protein at the ER–MT junctions.

Finally, we tested if tripartite-split-pFAST could be used to detect contacts/junctions between three organelles (Fig. 6j-l). Recent studies have suggested the existence of tri-organellar membrane contact sites, where three different organelles are closely juxtaposed^32^. A probe capable of specifically detecting such tri-organellar membrane contact sites has not been reported yet. As a proof-of-principle experiment, we used tripartite-split-pFAST to detect the formation of tri-organellar MT–ER–PM contact sites/junctions artificially induced using the split-FRB-rapamycin-FKBP CIT tool. We coexpressed ER-targeted pFAST_99-114_–^N^FRB–ECFP–Cb5, PM-targeted Lyn11–mCherry–^C^FRB–pFAST_1-98_ and MT-targeted TOM20–FKBP–pFAST_115-125_ in HeLa cells, and we treated cells with HBR-2,5DM. Addition of rapamycin led to a significant increase of the image correlation coefficient between the ER and PM signals (Fig. 6l), and the appearance of green fluorescence signal that colocalized with MT and patches between PM and ER (Fig. 6k and Supplementary Movie 8), in agreement with the assembly of tripartite-split-pFAST at the junction of the three organelles. Altogether, this set of experiments demonstrated the potential of tripartite-split-pFAST for the study of interactions at the contacts/junctions of organelles.

### Visualization of the Fos·Jun·DNA ternary complex

To further study the ability of tripartite-split-FAST to detect the proximity of three species, we next studied the Fos·Jun·DNA complex (Fig. 7). Fos and Jun interact together to form the heterodimeric transcription factor AP-1^33,34^. The dimerization of Fos and Jun is promoted by the interaction with DNA^35^. To visualize the Fos·Jun·DNA interaction, we fused the basic region-leucine zipper (bZIP) elements of the proteins Fos and Jun to the N-terminus of pFAST_115-125_ and pFAST_1-98_ respectively, and we indirectly targeted pFAST_99-114_ to DNA by fusing this third fragment at the C-terminus of the histone H2B involved in the structure of chromatin (Fig. 7a). HBR-2,5DM-treated HeLa cells co-expressing the three fusion proteins showed strong green fluorescence in the nucleus, in agreement with the assembly of the tripartite-split-FAST induced by the proximity of bFos, bJun and H2B (Fig. 7b,c). The selectivity of the complementation was demonstrated by using bFosΔ(179-193), a mutant of bFos lacking the residues 179-193, known to prevent the formation of heterodimers with bJun and the binding to DNA^34^. Accordingly, no fluorescence complementation was observed in HBR-2,5DM-treated HeLa cells co-expressing bFosΔ(179-193)-pFAST_115-125_, bJun-pFAST_1-98_ and H2B-pFAST_99-114_ (Fig. 7b,c), in agreement with the loss of the bFos·bJun·DNA complex. This set of experiments opens interesting prospects for studying the interaction of heterodimeric transcription factors with DNA or DNA-associated proteins, and further demonstrated the ability of tripartite-split-FAST to monitor the formation of ternary protein complexes and the identification of trimerization perturbation in cells.

**Figure 7.**
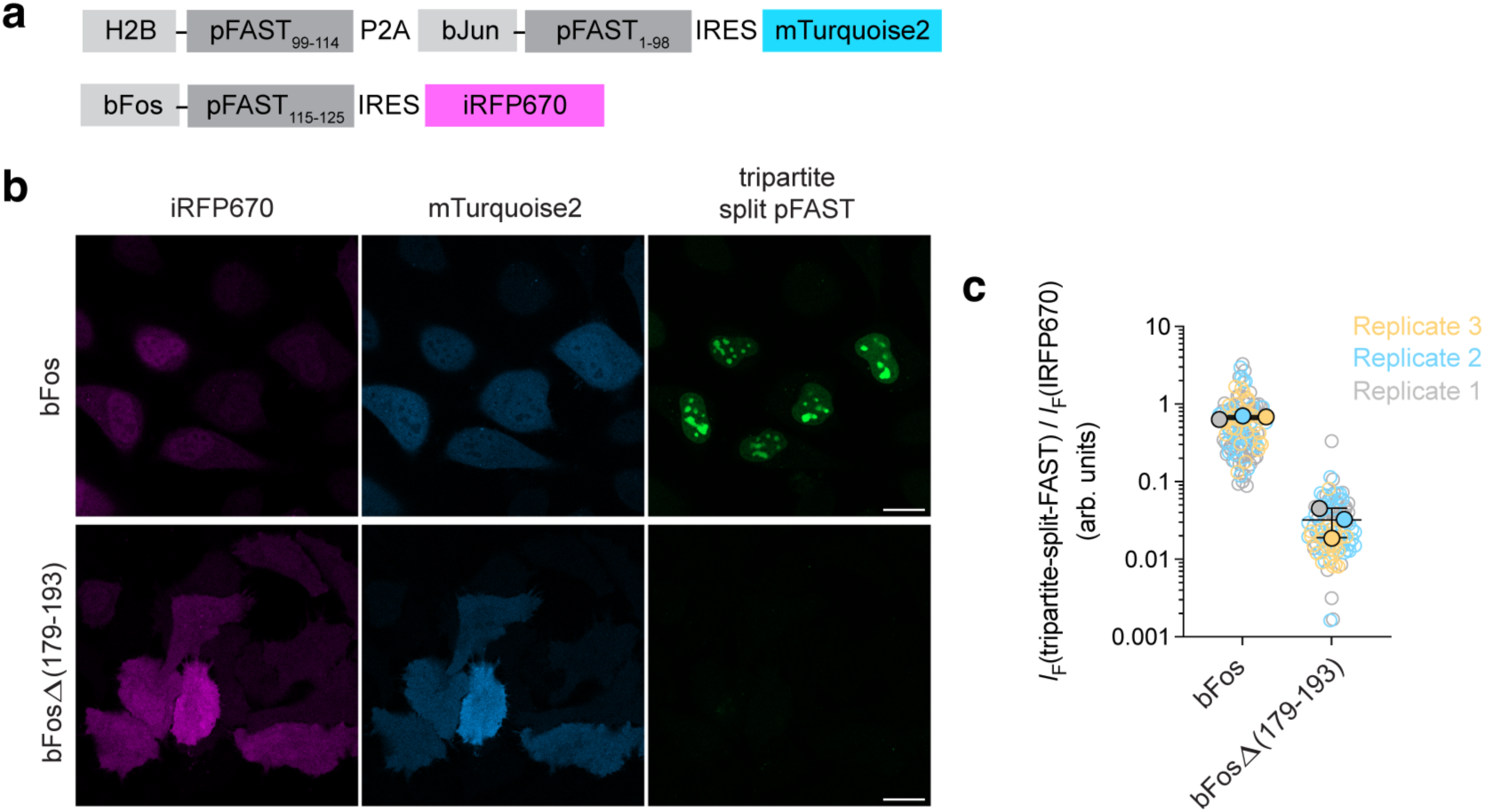
Visualization of the Fos·Jun·DNA ternary complex. **a** Constructs for the expression of bJun-pFAST_1-98_, bFos-pFAST_115-125_ and H2B-pFAST_99-114_ for probing the Fos·Jun·DNA ternary complex formation. **b** Representative confocal micrographs of HBR-2,5DM-treated HeLa cells co-expressing bJun-pFAST_1-98_, H2B-pFAST_99-114_ and either bFos-pFAST_115-125_ or bFosΔ(179-193)-pFAST_115-125_. Cells were treated with 10 μM HBR-2,5DM. Identical imaging settings were used for direct comparison. Scale bars 20 μm. Representative confocal micrographs of n > 100 cells from three independent experiments. **c** Relative nuclear tripartite-split-FAST fluorescence computed by normalizing the nuclear fluorescence intensity of tripartite-split-FAST *I*_F_(tripartite-split-FAST) with the fluorescence intensity *I*_F_(iRFP670) of co-expressed iRFP670. Each cell is color-coded according to the biological replicate it came from. The solid circles correspond to the mean of each biological replicate. The black line represents the mean ± SD of the three biological replicates.

## DISCUSSION

Most processes in cells are regulated through the specific assembly of multiprotein complexes. While FRET and BiFC have revolutionized the way one can visualize the proximity of two proteins, the study of ternary or higher order interactions remains highly challenging because very few molecular tools have been developed. To date, the proximity of three proteins can be visualized either (i) by fusing them to three fluorescent proteins that allow three mutually dependent energy transfer processes^18^, or (ii) by combining BiFC and FRET through fusions to a fluorescent protein acting as a FRET donor and two fragments of a split fluorescent protein acting as a FRET acceptor once in close proximity^19^.

In this study, we developed a reporter of ternary protein interactions based on the trisection of a variant of FAST, a small protein of 14 kDa that binds and stabilizes the fluorescent state of fluorogenic chromophores^12,13^. FAST and its variants were previously used for the design of reporters of binary interactions^8,11^. Bisection at position 114-115 generated two fragments that could efficiently complement into a functional reporter when in close proximity. These reporters display rapid and reversible complementation, allowing the monitoring of dynamic interactions with high temporal resolution. In the current study, we further showed that the cleavage of FAST or its variant ^RspA^FAST in between residues 98 and 99 generated two complementary fragments able to reassemble into functional reporters when in close proximity. Combination of the two split sites allowed us to generate a tripartite split system composed of three complementary fragments. We showed that efficient complementation requires the three fragments to be in close proximity to counteract the unfavorable entropy change associated to complementation, allowing thus the detection of ternary protein interactions. Optimal tripartite split system was obtained using pFAST, a FAST variant with higher conformational rigidity and higher fluorogen binding affinity^29^. We assume that the complementation of the three fragments is favored by a higher folding energy, and by the extra binding energy provided by fluorogen binding. We show that the complementation is rapid, allowing to monitor the formation of the ternary protein assembly within few minutes.

To our knowledge, only sfGFP and Nanoluciferase have been previously successfully split into three fragments capable of reassembling into a functional reporter^20,21^. Each system is composed of a large fragment, and two short fragments corresponding to consecutive beta strands, as in tripartite-split-pFAST. Complementation of tripartite-split-sfGFP can occur as soon as the two short fragments are in close proximity. This property was used to detect binary protein interactions with minimal fluorescence background. To do so, the two proteins are fused to the two short fragments. Upon formation of the binary protein complex, the large fragment spontaneously assembles with the two short fragments, leading to the reconstitution of a functional fluorescent protein within few hours (due to the slow maturation of the chromophore). Tripartite-split-Nanoluciferase displays similar behavior: reconstitution of functional Nanoluciferase can occur as soon as the two short fragments are in close proximity. This property allowed in a similar fashion to increase the signal-to background ratio of in vitro binary sandwich bioluminescent immunoassays^21,36^. To our knowledge, Tripartite-split-sfGFP and Tripartite-split-Nanoluciferase have however never been used for visualizing ternary protein complexes. Here, we present experiments showing the potential of tripartite split-pFAST for visualizing the close proximity of three proteins in the cytosol, at the plasma membrane and at the junction of different organelles with very high temporal resolution. Remarkably, unlike tripartite-split-sfGFP, the complementation kinetics in not limited by chromophore maturation, therefore tripartite-split-pFAST shows very rapid complementation, enabling to monitor in quasi real-time the formation of ternary protein assemblies by fluorescence microscopy.

Whether the approach can be general still needs however to be explored and demonstrated. We anticipate that the complementation of such tripartite split system will be more sensitive to geometrical constraints than bipartite split reporters. Appropriate geometry of the three fusions will be necessary to allow the reassembly of functional tripartite split-pFAST. To that end, the recent development of revolutionary protein structure prediction tools based on artificial intelligence^27^ should allow the optimization of the different domains’ position and orientation based on their structural features, overcoming the need for extensive experimental screening. In the future, we will focus our efforts on further exploring the potential of tripartite split-pFAST for various applications, including the visualization of proximity between three proteins, as well as the design of cellular biosensors. In particular, we believe that one field in which the tripartite-split-pFAST could find a broad interest is the field of membrane contact sites. Bipartite split-FAST was recently used to develop optical sensors enabling to monitor the dynamics of membrane contact sites^15^. Our results suggest that tripartite-split-pFAST could enable one to go one step further and study the recruitment of cytosolic molecular effectors at organelle membrane contacts/junctions with high spatial and temporal resolution, and could allow to detect contacts/junctions between three organelles, opening potentially new ways to explore the recently proposed existence of tri-organellar membrane contact sites^32^.

## METHODS

The presented research complies with all relevant ethical regulations.

### Statistics & Reproducibility

No sample size calculations were performed. When relevant, the sample size (n) is provided in the corresponding figure captions. Sample sizes were chosen to support meaningful conclusions. No data were excluded. The number of replicates for each individual experiments is indicated in the figure legends. All attempts at replication were successful. The experiments were not randomized. The investigators were not blinded to allocation during experiments and outcome assessment.

#### General

Synthetic oligonucleotides used for cloning were purchased from Integrated DNA technology. PCR reactions were performed with Q5 polymerase (New England Biolabs) in the buffer provided. PCR products were purified using QIAquick PCR purification kit (Qiagen). DNAse I, T4 ligase, Fusion polymerase, Taq ligase and Taq exonuclease were purchased from New England Biolabs and used with accompanying buffers and according to manufacturer protocols. Isothermal assemblies (Gibson assembly) were performed using homemade mix prepared according to previously described protocols^37^. Small-scale isolation of plasmid DNA was done using QIAprep miniprep kit (Qiagen) from 2 mL overnight bacterial culture supplemented with appropriate antibiotics. Large-scale isolation of plasmid DNA was done using the QIAprep maxiprep kit (Qiagen) from 150 mL of overnight bacterial culture supplemented with appropriate antibiotics. All plasmid sequences were confirmed by Sanger sequencing with appropriate sequencing primers (GATC Biotech). All the plasmids used in this study are listed in **Supplementary Table 1**, as well as DNA sequences of the open reading frames. Rapamycin was purchased from TCI. The synthesis of the fluorogens HMBR and HBR-2,5DM was previously reported^12,14^.

#### Cloning

The plasmids used in this study have been generated using isothermal Gibson assembly or restriction enzymes. The plasmid pAG398 allowing the expression of FAST-ENLYFQGGSGG-FAST-MYC was constructed by replacing the sequence coding for GGGSGGG by NLYFQGGSGG sequence, using primers containing the linker’s sequence for amplification, in the vector pAG328^38^ encoding FAST-GGGSGGG-FAST-MYC. The plasmid pAG404 allowing the bacterial expression of His-cpFAST(24-25)_ENLYFQGGSGG_ was constructed by inserting the sequence coding for FAST_25-125_-ENLYFQGGSGG-FAST_1-24_ sequence, amplified from the vector pAG398 encoding FAST-NLYFQGGSGG-FAST-MYC in the pET28 vector. Similarly, the plasmids pAG405, pAG406, pAG407, pAG408, pAG409 and pAG410 allowing the bacterial expression of His-cpFAST(35-36)_ENLYFQGGSGG_, His-cpFAST(59-60)_ENLYFQGGSGG_, His-cpFAST(72-73)_ENLYFQGGSGG_, His-cpFAST(89-90)_ENLYFQGGSGG_, His-cpFAST(98-99)_ENLYFQGGSGG_, and His-cpFAST(102-103)_ENLYFQGGSGG_, respectively, were constructed by inserting the sequence coding for CFAST_X-125_-ENLYFQGGSGG-FAST_1-Y_ sequence, amplified from the vector pAG398 encoding FAST-NLYFQGGSGG-FAST-MYC, in a pET28 vector. The plasmid pAG473 allowing the bacterial expression of His-cpFAST(24-25)_GGSGGSGGSGG_ was constructed by replacing the sequence coding for ENLYFQGGSGG by GGSGGSGGSGG sequence, using primers containing the linker’s sequence for amplification, in the vector pAG404 encoding His-cpFAST(24-25)_ENLYFQGGSGG_. Similarly, the plasmids pAG475, pAG478 pAG480 and pAG482 allowing the bacterial expression of His-cpFAST(24-25)_GGSGGSGG_, His-cpFAST(24-25)_GGSGG_, His-cpFAST(24-25)_GGS_, and His-cpFAST(24-25)_no linker_ respectively, were constructed by replacing the sequence coding for ENLYFQGGSGG by GGSGGSGG, GGSGG, GGS and no sequence, using primers containing the linker’s sequence for amplification, in the vector pAG404 encoding His-cpFAST(24-25)_ENLYFQGGSGG_. The plasmid pAG474 allowing the bacterial expression of His-cpFAST(98-99)_GGSGGSGGSGG_ was constructed by replacing the sequence coding for ENLYFQGGSGG by GGSGGSGGSGG sequence, using primers containing the linker’s sequence for amplification, in the vector pAG409 encoding His-cpFAST(98-99)_ENLYFQGGSGG_. Similarly, the plasmids pAG476, pAG479 pAG481 and pAG483 allowing the bacterial expression of His-cpFAST(98-99)_GGSGGSGG_, His-cpFAST(98-99)_GGSGG_, His-cpFAST(98-99)_GGS_, and His-cpFAST(98-99)_no linker_ respectively, were constructed by replacing the sequence coding for ENLYFQGGSGG by GGSGGSGG, GGSGG, GGS and no sequence, using primers containing the linker’s sequence for amplification, in the vector pAG409 encoding His-cpFAST(98-99)_ENLYFQGGSGG_. The plasmid pAG978 allowing bacterial expression of nanoFAST was constructed by replacing the sequence coding for FAST in the vector pAG087^12^ by the sequence coding for nanoFAST. The plasmid pAG935 allowing the mammalian expression of FAST_1-98_-FRB-MYC-IRES-HA-mTurquoise2 was constructed by replacing the sequence coding for FAST by FAST_1-98_ sequence, and inserting the sequence of FRB and the sequence of IRES-HA-mTurquoise2 amplified from the vector pAG573 encoding MYC-FRB-^RspA^FAST_1-114_-IRES-HA-mTurquoise2^11^, in the vector pAG104 encoding MYC-FAST^12^. The plasmid pAG802 allowing the mammalian expression of MYC-FKBP-FAST_99-125_-IRES-HA-iRFP670 was constructed by inserting the sequence coding for FAST_99-114_, using primers containing the insertion’s sequence for amplification, in the vector pAG577 encoding MYC-FKBP-FAST_115-125_-IRES-HA-iRFP670^11^. The plasmid pAG1095 allowing the mammalian expression of ^Rspa^FAST_1-98_-FRB-MYC-IRES-HA-mTurquoise2 was constructed by replacing the sequence coding for FAST_1-98_ by ^RspA^FAST_1-98_, amplified from the vector pAG573 encoding MYC-FRB-^RspA^FAST_1-114_-IRES-HA-mTurquoise2^11^, in the vector pAG935 encoding FAST_1-98_-FRB-MYC-IRES-HA-mTurquoise2. The plasmid pAG974 allowing the mammalian expression of MYC-FKBP-^Rspa^FAST_99-125_-IRES-HA-iRFP670 by replacing the sequence coding for FAST_99-125_ by ^RspA^FAST_99-125_, amplified from the vector pAG580 encoding MYC-FKBP-^RspA^FAST_115-125_-IRES-HA-iRFP670^11^, in the vector pAG802 encoding MYC-FKBP-FAST_99-125_-IRES-HA-iRFP670. The plasmid pAG1127 allowing the mammalian expression of MYC-^RspA^FAST_99-114_-FRB-^RspA^FAST_1-98_-IRES-HA-mTurquoise2 was constructed by replacing the sequence coding for FAST1-114 (a.k.a. NFAST) by ^RspA^FAST_1-98_ sequence, amplified from the vector pAG573^11^, and inserting the sequence of ^RspA^FAST_99-114_, amplified from the vector pAG573, in the vector pAG490^8^ encoding FRB-Hha(N)-IRES-mTurquoise2. The plasmid pAG580 allowing the mammalian expression of MYC-FKBP-^RspA^FAST_115-125_-IRES-HA-IRFP670 was previously described^11^. The plasmid pAG1148 allowing the mammalian expression of MYC-pFAST_99-114_-FRB-pFAST_1-98_-IRES-HA-mTurquoise2 was constructed by replacing the sequences coding for ^RspA^FAST_99-114_ and ^RspA^FAST_1-98_ by the sequences encoding for pFAST_99-114_ and pFAST_1-98_ respectively, both amplified from the vector pAG654^29^, in the vector pAG1127. The plasmid pAG1151 allowing the mammalian expression of MYC-FKBP-pFAST_115-125_-IRES-HA-IRFP670 was constructed by replacing the sequence coding for ^RspA^FAST_115-125_ by the pFAST_115-125_ sequence, amplified from the vector pAG654^29^, in the vector pAG580^11^ encoding FKBP-^RspA^FAST_115-125_-IRES-IRFP670. The plasmid pAG1184 allowing the mammalian expression of MYC-pFAST_99-114-_^N^FRB-P2A-^C^FRB-pFAST_1-98_-IRES-HA-mTurquoise2 was constructed by using primers containing the P2A sequence for the insertion at the nucleotide position 114/115 of the FRB sequence of the vector pAG1148 encoding MYC-pFAST_99-114_-FRB-pFAST_1-98_-IRES-HA-mTurquoise2. The plasmid pAG1223 allowing the mammalian expression of Lyn11-mCherry-^C^FRB-pFAST_1-98_ was constructed by replacing the PYP sequence by the sequence encoding for ^C^FRB-pFAST_1-98_, amplified from pAG1184 and introducing the Lyn11 sequence, by using primers containing the sequence, in the vector pAG097^12^ encoding for mCherry-PYP. The plasmid pAG1224 allowing the mammalian expression of MYC-pFAST_99-114_-^N^FRB-ECFP-Cb5 was generated by replacing the sequence coding for P2A-^C^FRB-pFAST_1-98_-IRES-HA-mTurquoise2 by the ECFP-Cb5 sequence, amplified from pAG1169^28^ encoding ^FIRE^mate-ECFP-Cb5, in the vector pAG1184. The plasmid pAG1239 allowing the mammalian expression MYC-pFAST_99-114_-^N^FRB-ECFP-Caax was generated by replacing the sequence encoding Cb5 by the Caax sequence, by using primers containing the sequence, in the vector pAG1224. The plasmid pAG1240 allowing the mammalian expression of TOM20-mCherry-^C^FRB-pFAST_1-98_ was constructed by replacing the Lyn11 sequence by TOM20 sequence, amplified from the Addgene vector #171461 Tom20-CR coding for TOM20-ECFP-FRB^39^, in the vector pAG1223. The plasmid pAG1225bis allowing the mammalian expression of TOM20-FKBP-pFAST_115-125_ was generated by replacing the sequence encoding ECFP-FRB by the FKBP-pFAST_115-125_ sequence, amplified from pAG1151, in the Addgene vector #171461 Tom20-CR coding for TOM20-ECFP-FRB^39^. The plasmid pAG1713 allowing the mammalian expression of H2B-pFAST_99-114_-P2A-bJun-pFAST_1-98_-IRES-HA-mTurquoise2 was constructed by adding H2B sequence to pAG1256 (CMV pFAST_99-114_-P2A-bJun-pFAST_1-98_-IRES-mTurquoise2, which was itself constructed by adding bJun sequence from Addgene vector #22012 pBiFC-bJunVN173 to pAG1184) for expression of a fusion between H2B and pFAST_99-114_. The plasmid pAG1258 allowing the mammalian expression of myc-bFos-pFAST_115-125_-IRES-HA-iRFP670 was constructed by replacing FKBP sequence from pAG1151^28^ by bFos sequence from Addgene vector #22013 pBiFC-bFosVC155. The plasmid pAG1265 allowing the mammalian expression of myc-bFosΔ(179-193)-pFAST_115-125_-IRES-HA-iRFP670 was constructed by replacing FKBP sequence from pAG1151^28^ by bFosΔ(179-193) sequence from Addgene vector #22014 pBiFC-bFosDeltaZipVC155.

#### Protein expression in bacteria and purification

Plasmids encoding the circular permutations or NanoFAST fused to an His-tag were transformed in BL21 (DE3) competent *E. coli* (New England Biolabs). Cells were grown at 37 °C in lysogen Broth (LB) medium supplemented with 50 μg/ml kanamycin to OD_600nm_ 0.6. Expression was induced overnight at 16 °C by adding isopropyl-β-D-1-thiogalactopyranoside (IPTG) to a final concentration of 1 mM. Cells were harvested by centrifugation (4300 × g for 20 min at 4 °C) and frozen. For purification, the cell pellet was resuspended in lysis buffer (PBS supplemented with 2.5 mM MgCl_2_, 1 mM of protease inhibitor PhenylMethaneSulfonyl Fluoride PMSF, 0.025 mg/mL of DNAse, pH 7.4) and sonicated (5 min, 20% of amplitude) on ice. The lysate was incubated for 2 h on ice to allow DNA digestion by DNAse. Cellular fragments were removed by centrifugation (9000 × g for 1 h at 4 °C). The supernatant was incubated overnight at 4 °C by gentle agitation with pre-washed Ni-NTA agarose beads in PBS buffer complemented with 20 mM of imidazole. Beads were washed with 10 volumes of PBS complemented with 20 mM of imidazole and with 5 volumes of PBS complemented with 40 mM of imidazole. His-tagged proteins were eluted with 5 volumes of PBS complemented with 0.5 M of imidazole. The buffer was exchanged to PBS (0.05 M phosphate buffer, 0.150 M NaCl) using PD-10 desalting columns (GE Healthcare). Purity of the proteins were evaluated using SDS-PAGE electrophoresis stained with Coomassie blue.

#### Physico-chemical measurements

Steady state UV-Vis and fluorescence spectra were recorded at 25°C on a Spark 10M (Tecan). Fluorescence quantum yield measurements were determined as previously described^12^ using FAST:HMBR as a reference. Reciprocal dilution with protein solution was used so as to keep the protein concentration constant at 40 μM while diluting the fluorogen solution. Thermodynamic dissociation constants were determined by titration experiments in which we measured the fluorescence of the fluorescent assembly at various fluorogen concentrations using a Spark 10 M plate reader (Tecan) and fitting data to a one-site specific binding model. Data were processed using GraphPad Prism v9.3.0.

#### Cell culture

HeLa cells (ATCC CRM-CCL2) were cultured in Minimal Essential Media (MEM) supplemented with phenol red, Glutamax I, 1 mM of sodium pyruvate, 1% (vol/vol) of non-essential amino-acids and 10% (vol/vol) fetal calf serum (FCS), at 37 °C in a 5% CO_2_ atmosphere. HEK293T (ATCC CRL-3216) cells were cultured in Dulbecco’s Modified Eagle Medium (DMEM) supplemented with phenol red and 10% (vol/vol) fetal calf serum at 37 °C in a 5% CO_2_ atmosphere. U2OS cells (ATCC HTB-96) were cultured in Dulbecco’s modified Eagle’s medium (DMEM) supplemented with phenol red and 10% (vol/vol) fetal calf serum and 1% (vol/vol) penicillin–streptomycin at 37°C in a 5% CO_2_ atmosphere. For imaging, cells were seeded in μDish IBIDI (Biovalley) coated with poly-L-lysine. Cells were transiently transfected using Genejuice (Merck) according to the manufacturer’s protocols for 24 h prior to imaging. Cells were washed with DPBS (Dulbecco’s Phosphate-Buffered Saline), and treated with DMEM media (without serum and phenol red) supplemented with the fluorogen at the indicated concentration few minutes prior to imaging. To follow the kinetics of rapamycin-induced complementation, rapamycin was added at t = 0 min.

#### Flow cytometry analysis

Flow cytometry in HEK293T cells was performed on a MACSQuant® analyzer equipped with 405 nm, 488 nm and 635 nm lasers and seven channels. To prepare samples, cells were first grown in cell culture flasks, then transiently co-transfected 24 hours after seeding using Genejuice (Merck) according to the manufacturer’s protocol for 24 hours. After 24 hours, cells were centrifuged in PBS with BSA (1 mg/ml) and resuspend in PBS-BSA supplemented with the appropriate amounts of compounds. For each experiment, 20,000 cells positively expressing mTurquoise2 (Ex 434 / Em 450 ± 25) and iRFP670 (Ex 638 nm / Em 660 ± 10 nm) were analyzed with the following parameters: Ex 488nm, Em 525 ± 20 nm. Data were analyzed using FlowJo v10.7.1. A figure presenting the gating strategy is presented in supporting information on **Supplementary Fig. 6**. The analysis of the complementation was performed as follows: the evolution of the mean cell fluorescence in the function of the concentration of fluorogen in the presence and in the absence of rapamycin was fitted with a one-site binding model equation with GraphPad Prism v9.3.0, sharing the maximal value for the experiments in the presence and in the absence of rapamycin. This allows us to determine the effective concentration of fluorogen for half-maximal complementation (EC_50_) as well as the theoretical maximum fluorescence value. This latter value is then used to compute the percentage of complementation for each concentration of fluorogen.

#### Live-cell fluorescence microscopy

Confocal micrographs were acquired on a Zeiss LSM 980 Laser Scanning Microscope equipped with a plan apochromat 63× /1.4 NA oil immersion objective. ZEN software was used to collect the data. The images were analyzed using Icy (2.4.0.0) or Fiji (Image J). To track fluorescence signal in a specific organelle, the organelle contour was determined by masking the signal in the ECFP channel with the plug-in *HK-means* with the following parameters: intensity class equals 100, min object size (px) equals 500, max object size (px) equals 1500-3000. The signal intensity of the ROI was tracked over the time for each channel by using plug-in *Active Contours.* The background signal was subtracted. Data were processed using GraphPad Prism v9.3.0. Formation of organelle junctions was assessed computing image correlation coefficient using Fisher’s transform of Pearson’s correlation coefficient as previously reported^28^. In brief, to determine the Pearson’s correlation between two channels, we selected three ROI of 50×50 pixels for each cell and we used the Coloc2 plugin on FIJI. For the analysis of the trimerization the Pearson’s correlation was calculated before and after the addition of rapamycin. Fisher’s transform was applied to normalize Pearson’s correlation coefficient.

## DATA AVAILABILITY

Data supporting the findings of this study are available within the article and supplementary information, and are available from the corresponding authors upon request. Requests will be fulfilled within four weeks. Source data are provided with this paper. The plasmids developed in this study may be requested from the corresponding author.

## ACKNOWLEDGMENTS

We thank the imaging facility of the Institut de Biologie Paris Seine of Sorbonne University. We thank Takafumi Miyamoto for the plasmid Tom20-CR (Addgene plasmid # 171461). This work has been supported by the European Research Council (ERC-2016-CoG-724705 FLUOSWITCH), the Institut Universitaire de France and the Dynamic Imaging program of the Chan-Zuckerberg Initiative DAF (grant number 2023-321185), an advised fund of Silicon Valley Community.

## AUTHOR CONTRIBUTION STATEMENT

S.B., F.B, A.Ge., A.B., L.E.H, H.B. and A.Ga. designed the experiments. S.B., F.B, A.Ge., A.B, L.E.H. and H.B. performed the experiments. S. B., F.B, A.Ge., A.B., L.E.H, H.B. and A.Ga. analyzed the experiments. S. B., F.B, and A.Ga. wrote the paper with the help of the other authors.

