## Supplementary Information for "A tripartite chemogenetic fluorescent reporter: design and evaluation for visualizing ternary protein complexes"

**This PDF file includes:**

Legends of Supplementary Movies

Supplementary Figs 1-9

Supplementary Table 1

Supplementary references

### Legends of Supplementary Movies

**Supplementary Movie 1.** HEK293T cells expressing $^{RspA}FAST_{1\text{-}98}$-FRB and FKBP-$^{RspA}FAST_{99\text{-}125}$ were treated with 10 μM HMBR. Cells were imaged by time-lapse confocal microscopy after addition of 500 nM of rapamycin. Experiments were repeated two times with similar results. Scale bar 20 μm.

**Supplementary Movie 2.** HEK293T cells expressing $pFAST_{99\text{-}114}$–FRB–$pFAST_{1\text{-}98}$ and FKBP–$pFAST_{115\text{-}125}$ were treated with 5 μM HBR-2,5DM. Cells were imaged by time-lapse confocal microscopy after addition of 500 nM of rapamycin. Experiments were repeated three times with similar results. Scale bar 20 μm.

**Supplementary Movie 3.** HEK293T cells expressing the triplet $pFAST_{99\text{-}114}$–$^{N}$FRB, $^{C}$FRB–$pFAST_{1\text{-}98}$ and FKBP–$pFAST_{115\text{-}125}$ were treated with 10 μM HBR-2,5DM. Cells were imaged by time-lapse confocal microscopy after addition of 500 nM of rapamycin. Experiments were repeated three times with similar results. Scale bar 20 μm.

**Supplementary Movie 4.** HEK293T cells expressing the triplet $pFAST_{99\text{-}114}$–$^{N}$FRB–ECFP–CAAX, Lyn11-mCherry–$^{C}$FRB–$pFAST_{1\text{-}98}$ and FKBP–$pFAST_{115\text{-}125}$ were treated with 10 μM HBR-2,5DM. Cells were imaged by time-lapse confocal microscopy after addition of 500 nM of rapamycin. Experiments were repeated three times with similar results. Scale bar 20 μm.

**Supplementary Movie 5.** HeLa cells co-expressing $pFAST_{99\text{-}114}$–$^{N}$FRB–ECFP–CAAX, Tom20-mCherry–$^{C}$FRB–$pFAST_{1\text{-}98}$ and FKBP–$pFAST_{115\text{-}125}$ were treated with 10 μM HBR-2,5DM. Cells were imaged by time-lapse confocal microscopy after addition of 500 nM of rapamycin. Experiments were repeated three times with similar results. Scale bar 20 μm.

**Supplementary Movie 6.** HeLa cells co-expressing $pFAST_{99\text{-}114}$–$^{N}$FRB–ECFP–Cb5, Lyn11-mCherry–$^{C}$FRB–$pFAST_{1\text{-}98}$ and FKBP–$pFAST_{115\text{-}125}$ were treated with 10 μM HBR-2,5DM. Cells were imaged by time-lapse confocal microscopy after addition of 500 nM of rapamycin. Experiments were repeated three times with similar results. Scale bar 20 μm.

**Supplementary Movie 7.** HeLa cells co-expressing $pFAST_{99\text{-}114}$–$^{N}$FRB–ECFP–Cb5, Tom20-mCherry–$^{C}$FRB–$pFAST_{1\text{-}98}$ and FKBP–$pFAST_{115\text{-}125}$ were treated with 10 μM HBR-2,5DM. Cells were imaged by time-lapse confocal microscopy after addition of 500 nM of rapamycin. Experiments were repeated three times with similar results. **h** Representative micrographs before and after addition of rapamycin. Scale bar 20 μm.

**Supplementary Movie 8.** HeLa cells co-expressing $pFAST_{99\text{-}114}$–$^{N}$FRB–ECFP–Cb5, Lyn11-mCherry–$^{C}$FRB–$pFAST_{1\text{-}98}$, and Tom20–FKBP–$pFAST_{115\text{-}125}$ were treated with 10 μM HBR-2,5DM. Cells were imaged by time-lapse confocal microscopy after addition of 500 nM of rapamycin. Experiments were repeated three times with similar results. Scale bar 20 μm.

**a**

| Position | Residue | Estimated probability | Position | Residue | Estimated probability | Position | Residue | Estimated probability |
|---|---|---|---|---|---|---|---|---|
| 1 | M | 0.822 | 43 | N | 0.073 | 85 | S | 0.676 |
| 2 | E | 0.822 | 44 | A | 0.044 | 86 | G | 0.689 |
| 3 | H | 0.858 | 45 | A | 0.020 | 87 | N | 0.836 |
| 4 | V | 0.833 | 46 | E | 0.000 | 88 | L | 0.793 |
| 5 | A | 0.584 | 47 | G | 0.018 | 89 | N | 0.936 |
| 6 | F | 0.429 | 48 | D | 0.087 | 90 | T | 0.853 |
| 7 | G | 0.491 | 49 | I | 0.133 | 91 | M | 0.411 |
| 8 | S | 0.878 | 50 | T | 0.536 | 92 | F | 0.462 |
| 9 | E | 0.942 | 51 | G | 0.824 | 93 | E | 0.289 |
| 10 | D | 0.951 | 52 | R | 0.960 | 94 | Y | 0.249 |
| 11 | I | 0.420 | 53 | D | 0.976 | 95 | T | 0.193 |
| 12 | E | 0.067 | 54 | P | 0.271 | 96 | F | 0.498 |
| 13 | N | 0.336 | 55 | K | 0.442 | 97 | D | 0.662 |
| 14 | T | 0.547 | 56 | Q | 0.353 | 98 | Y | 0.909 |
| 15 | L | 0.609 | 57 | V | 0.611 | 99 | Q | 0.840 |
| 16 | A | 0.658 | 58 | I | 0.311 | 100 | M | 0.811 |
| 17 | K | 0.336 | 59 | G | 0.689 | 101 | T | 0.902 |
| 18 | M | 0.758 | 60 | K | 0.707 | 102 | P | 0.971 |
| 19 | D | 0.838 | 61 | N | 0.093 | 103 | T | 0.936 |
| 20 | D | 0.653 | 62 | F | 0.004 | 104 | K | 0.622 |
| 21 | G | 0.460 | 63 | F | 0.036 | 105 | V | 0.160 |
| 22 | Q | 0.122 | 64 | K | 0.182 | 106 | K | 0.049 |
| 23 | L | 0.111 | 65 | D | 0.200 | 107 | V | 0.016 |
| 24 | D | 0.724 | 66 | V | 0.160 | 108 | H | 0.073 |
| 25 | G | 0.796 | 67 | A | 0.064 | 109 | M | 0.007 |
| 26 | L | 0.358 | 68 | P | 0.247 | 110 | K | 0.024 |
| 27 | A | 0.307 | 69 | C | 0.562 | 111 | K | 0.031 |
| 28 | F | 0.213 | 70 | T | 0.538 | 112 | A | 0.191 |
| 29 | G | 0.036 | 71 | D | 0.838 | 113 | L | 0.653 |
| 30 | A | 0.007 | 72 | S | 0.951 | 114 | S | 0.833 |
| 31 | I | 0.007 | 73 | P | 0.936 | 115 | G | 0.924 |
| 32 | Q | 0.027 | 74 | E | 0.882 | 116 | D | 0.951 |
| 33 | L | 0.018 | 75 | F | 0.451 | 117 | S | 0.733 |
| 34 | D | 0.458 | 76 | Y | 0.073 | 118 | Y | 0.227 |
| 35 | G | 0.953 | 77 | G | 0.200 | 119 | W | 0.053 |
| 36 | D | 0.971 | 78 | K | 0.140 | 120 | V | 0.073 |
| 37 | G | 0.682 | 79 | F | 0.153 | 121 | F | 0.033 |
| 38 | N | 0.413 | 80 | K | 0.256 | 122 | V | 0.038 |
| 39 | I | 0.038 | 81 | E | 0.356 | 123 | K | 0.136 |
| 40 | L | 0.116 | 82 | G | 0.298 | 124 | R | 0.598 |
| 41 | Q | 0.027 | 83 | V | 0.178 | 125 | V | 0.820 |
| 42 | Y | 0.042 | 84 | A | 0.400 | | | |

**b**

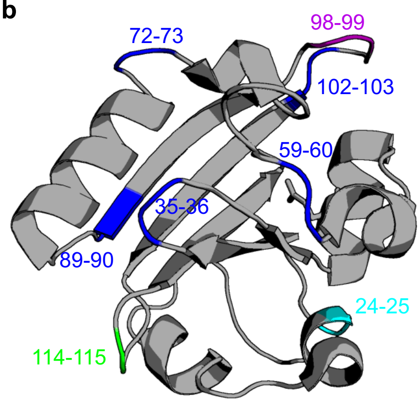


**Supplementary Figure 1. Identification of the split sites for the design of putative circular permutations of FAST. a** Probability for the circular permutation of the photoactive yellow protein PYP (PDB 1NWZ), the parental protein of FAST, estimated using CPred. **b** Split sites retained for the generation of circular permutations of FAST and further experimental analysis.

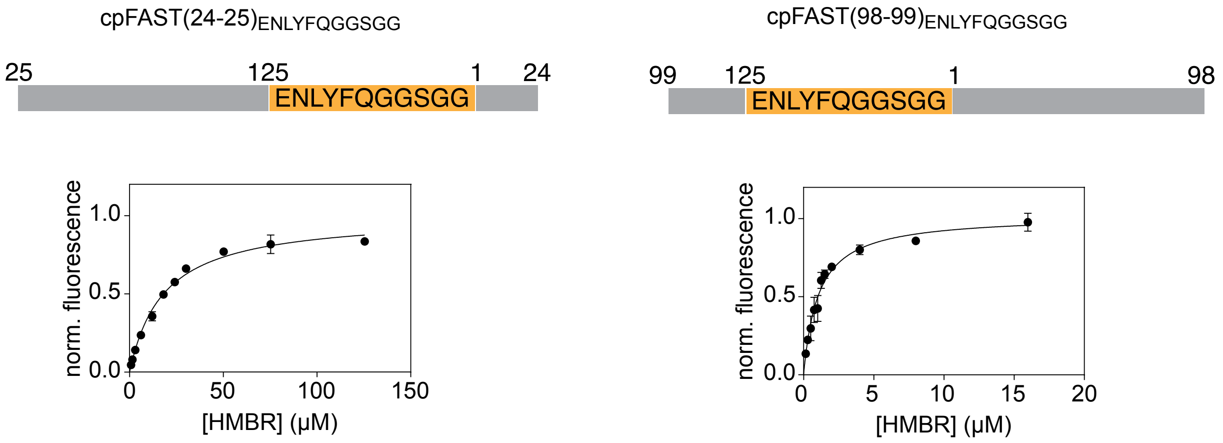


**Supplementary Figure 2. Characterization of the fluorogen binding affinity of cpFAST(24-25)$_{\mathbf{ENLYFQGGSGG}}$ and cpFAST(98-99)$_{\mathbf{ENLYFQGGSGG}}$.** HMBR titration curves in pH 7.4 HEPES buffer (50 mM) containing NaCl (150 mM). The protein concentration was fixed to 0.1 μM. Data represent the mean ± SD of three experiments. Least-squares fit (line) gave the thermodynamic dissociation constant $K_{D,HMBR}$ provided in **Fig. 2a**.

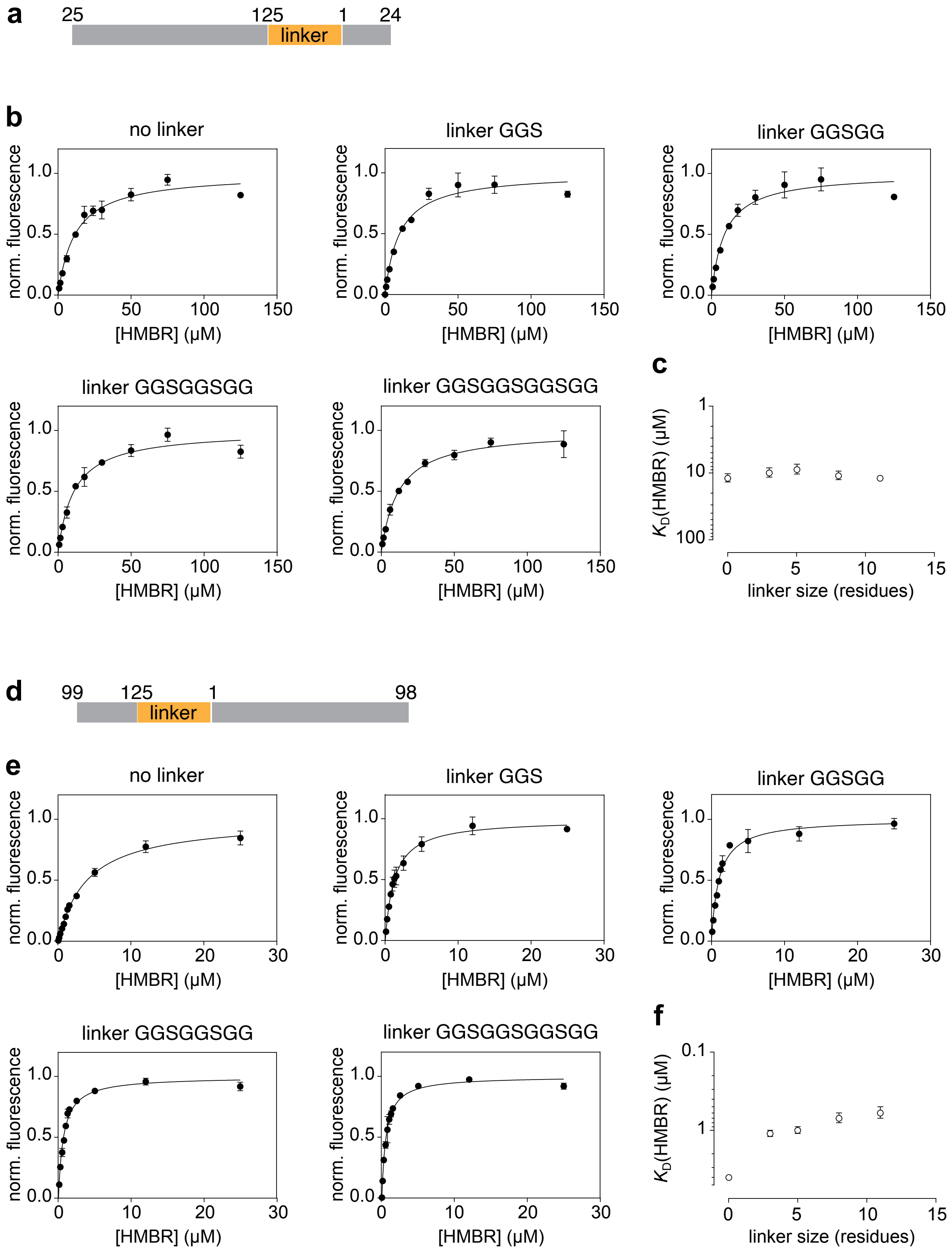


**Supplementary Figure 3. Influence of the linker in cpFAST(24-25) and cpFAST(98-99). a,d** Schematic representation of cpFAST(24-25)$_{linker}$ and cpFAST(98-99)$_{linker}$, respectively**. b,e** HMBR titration curves for the different circular permutations, in pH 7.4 HEPES buffer (50 mM) containing NaCl (150 mM). The protein concentration was fixed to 0.1 µM. Data represent the mean ± SD of three experiments. Least-squares fit (line) gave the thermodynamic dissociation constant $K_{D,HMBR}$ provided in **Fig. 2b**. **c,f** Evolution of HMBR dissociation constants in function of the linker size.

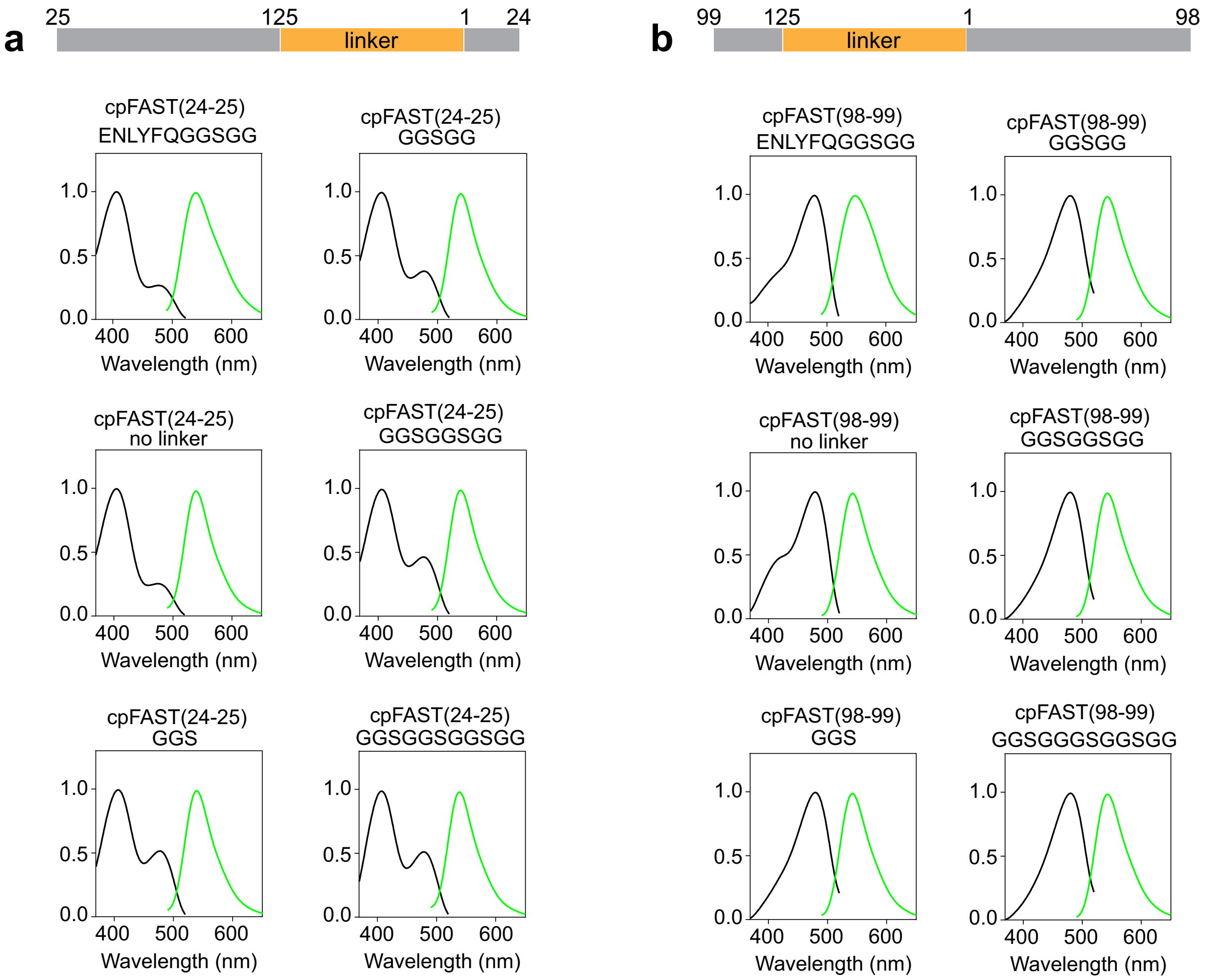


**Supplementary Figure 4. Spectral properties of the cpFAST:HMBR assemblies.** Normalized absorption (black) and emission (green) spectra of the cpFAST(24-25)$_{\text{linker}}$ (**a**) and cpFAST(98-99)$_{\text{linker}}$ (**b**) with HMBR. Spectra for ENLYFQGGSGG and other GGS linkers were recorded using 3 μM and 1.5 μM HMBR, respectively, and 40 μM protein in pH 7.4 phosphate buffer saline at 25°C.

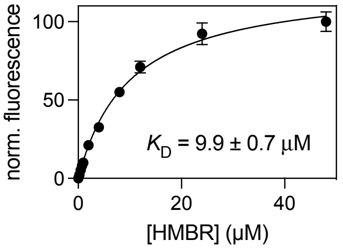


**Supplementary Figure 5. Characterization of the fluorogen binding affinity of nanoFAST.** HMBR titration curves for nanoFAST, in pH 7.4 phosphate buffer (50 mM) containing NaCl (150 mM). The protein concentration was fixed to 0.5 µM. Data represent the mean ± SD of three experiments. Least-squares fit (line) gave the indicated thermodynamic dissociation constant $K_D$.

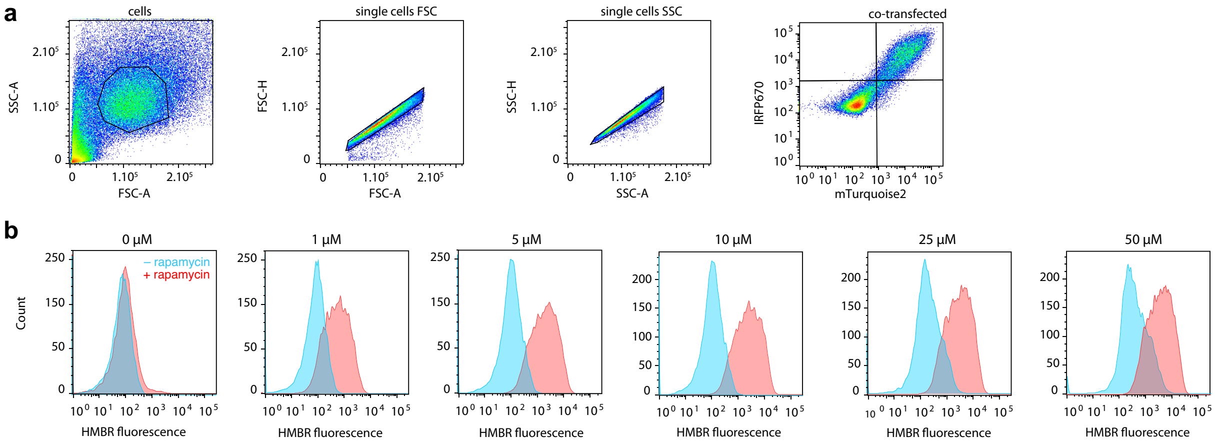


**Supplementary Figure 6. Flow cytometry analysis. a** Gating strategy. Side versus forward scatter (SSC-A vs FSC-A) density plot was used to identify cells based on size and granularity. Secondly, forward scatter height (FSC-H) vs. forward scatter area (FSC-A) density plot and then side scatter height (SSC-H) vs side scatter area (SSC- A) plot were used to select single cells. Co-transfected cells were then selected by using the signal of IRFP670 and mTurquoise2 transfection reporters. **b** Representative fluorescence analysis. HEK293T cells co-expressing the FK506-binding protein (FKBP) fused at the N-terminus of $^{RspA}$FAST$_{99-125}$ and the FKBP-rapamycin-binding domain of mammalian target of rapamycin (FRB) fused to the C-terminus of $^{RspA}$FAST$_{1-98}$ were treated without (blue) or with (red) 500 nM of rapamycin, and with 1, 5, 10, 25 or 50 µM of HMBR and then analyzed by flow cytometry.

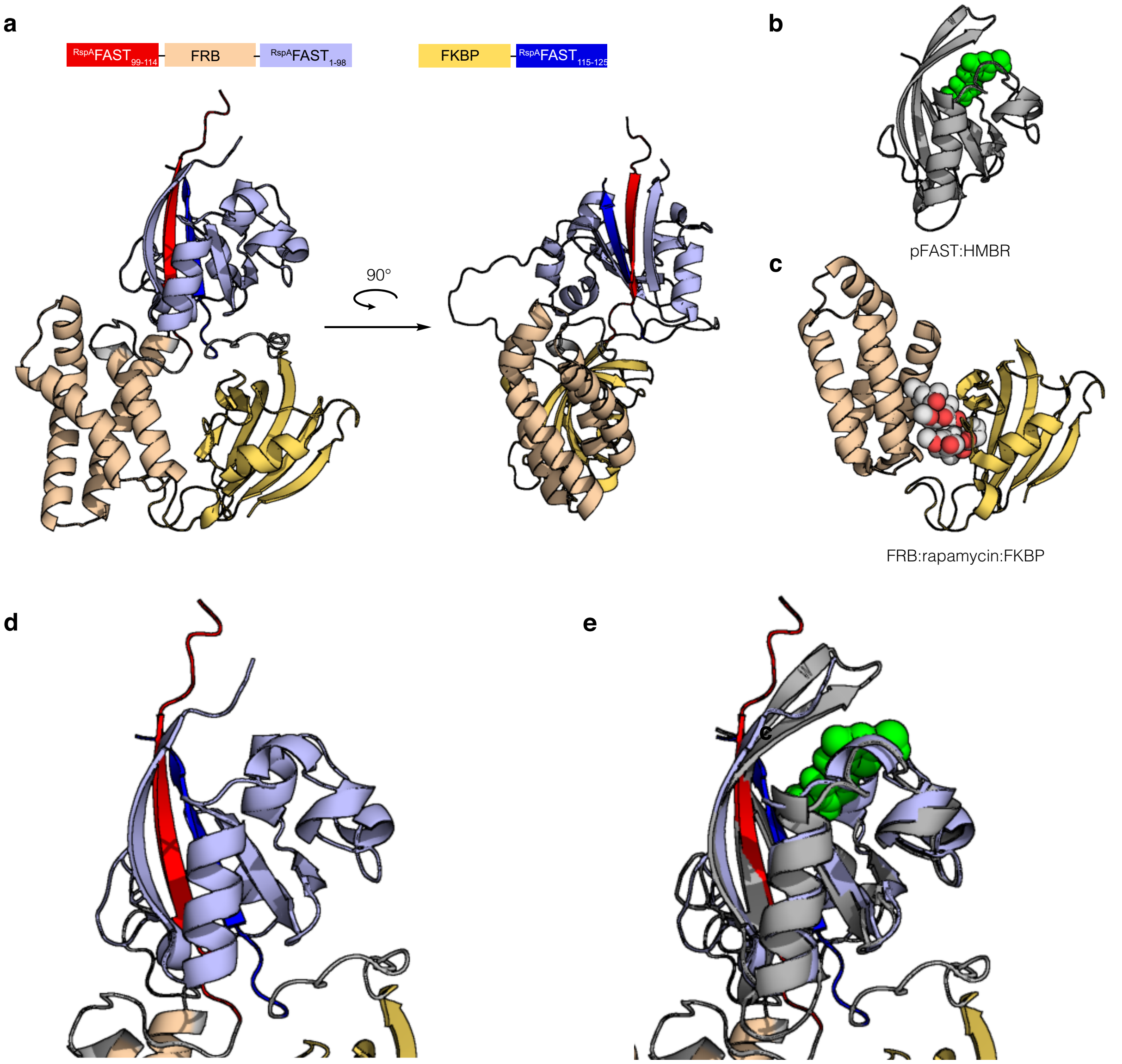


**Supplementary figure 7. Alphafold-assisted topology choice. a** Alphafold[1] prediction of the $^{RspA}FAST_{99\text{-}114}$–FRB–$^{RspA}FAST_{1\text{-}98}$ / FKBP–$^{RspA}FAST_{115\text{-}125}$ interaction. **b** Model of the assembly pFAST:HMBR (ref. 2). **c** Structure of the FRB:rapamycin:FKBP complex (PDB 3FAP). **d** zoom on the assembled tripartite RspA-splitFAST. **e** zoom on the assembled tripartite RspA-splitFAST with superimposed pFAST:HMBR (pFAST in grey, HMBR in green).

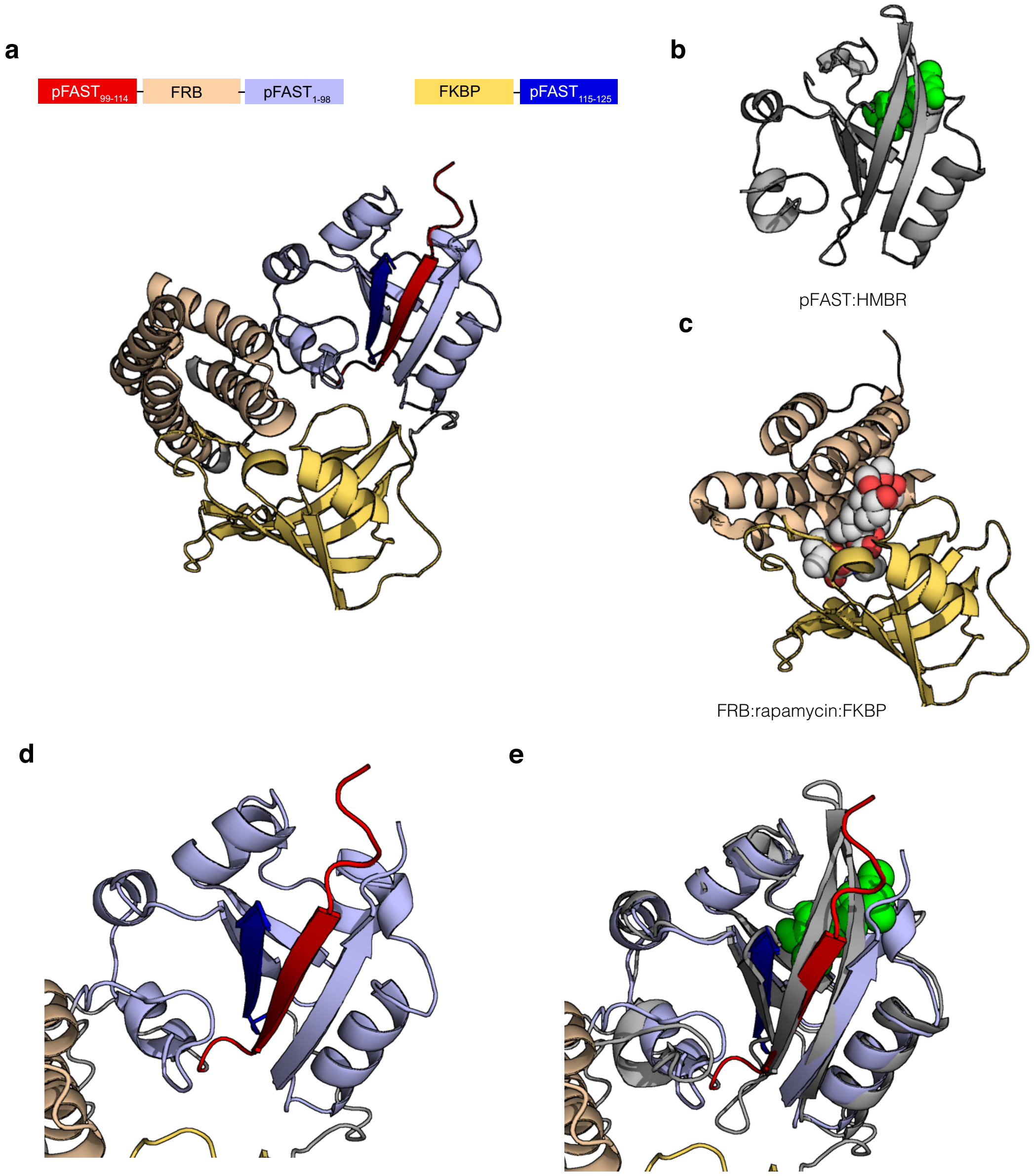


**Supplementary figure 8. Alphafold-assisted topology choice. a** Alphafold[1] prediction of the pFAST$_{99-114}$–FRB–pFAST$_{1-98}$ / FKBP–pFAST$_{115-125}$ interaction. **b** Model of the assembly pFAST:HMBR (ref. 2). **c** Structure of the FRB:rapamycin:FKBP complex (PDB 3FAP). **d** zoom on the assembled tripartite split pFAST. **e** zoom on the assembled tripartite split pFAST with superimposed pFAST:HMBR (pFAST in grey, HMBR in green).

**a**

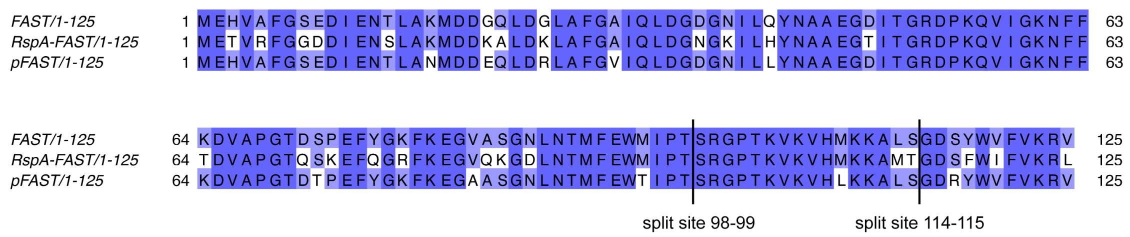


**b**

| | FAST | RspA-FAST | pFAST |
|---|---|---|---|
| **full-length** | $K_{D,HMBR}$ = 0.13 μM (see ref 3) | $K_{D,HMBR}$ = 0.03 μM (see ref 4) | $K_{D,HMBR}$ = 0.01 μM (see ref 3) |
| **bipartite 114-115** | $EC_{50,–interaction}$ (HMBR) = 18 μM (see ref 4)<br>$EC_{50,+interaction}$ (HMBR) = 3.1 μM (see ref 4) | $EC_{50,–interaction}$ (HMBR) = 100 μM (see ref 4)<br>$EC_{50,+interaction}$ (HMBR) = 1.0 μM (see ref 4) | $EC_{50,–interaction}$ (HMBR) = 0.5 μM (see ref 5)<br>$EC_{50,+interaction}$ (HMBR) = 0.5 μM (see ref 5) |
| **bipartite 98-99**<br>(See Fig. 3) | $EC_{50,–interaction}$ (HMBR) = 31 μM<br>$EC_{50,+interaction}$ (HMBR) = 7.5 μM | $EC_{50,–interaction}$ (HMBR) = 270 μM<br>$EC_{50,+interaction}$ (HMBR) = 3.0 μM | |
| **tripartite**<br>(with 2 proteins)<br>(See Fig. 4) | | $EC_{50,–interaction}$ (HMBR) > 500 μM<br>$EC_{50,+interaction}$ (HMBR) = 123 μM | $EC_{50,–interaction}$ (HMBR) > 500 μM<br>$EC_{50,+interaction}$ (HMBR) = 12 μM |
| | | $EC_{50,–interaction}$ (HBR-2,5DM) > 500 μM<br>$EC_{50,+interaction}$ (HBR-2,5DM) = 8.1 μM | $EC_{50,–interaction}$ (HBR-2,5DM) = 57 μM<br>$EC_{50,+interaction}$ (HBR-2,5DM) = 1.2 μM |
| **tripartite**<br>(with 3 proteins)<br>(See Fig. 5) | | | $EC_{50,–interaction}$ (HBR-2,5DM) = 91 μM<br>$EC_{50,+interaction}$ (HBR-2,5DM) = 0.31 μM |

**Supplementary Figure 9. FAST variants and their bipartite and tripartite versions. a** Alignment of the sequences of FAST, RspA-FAST and pFAST with the split sites 98-99 and 114-115 indicated**. b** Summary of the properties of their full-length, bipartite 114-115, bipartite 98-99 and tripartite versions. For the full-length proteins, the thermodynamic dissociation constants of the complex with the fluorogen HMBR ($K_{D,HMBR}$) are provided. For the bipartite and tripartite systems, the effective concentration for half maximal complementation in cells is given, either in the absence ($EC_{50,–interaction}$) or the presence ($EC_{50,+interaction}$) of an interaction. The fluorogens used are specified in brackets.

### Supplementary Table 1. Plasmids used in this study

| Plasmid | ORF | ORF sequence |
|---|---|---|
| pAG398 | FAST-ENLYFQGGSGG-FAST-MYC | atggaacatgtcgccttcggcagcgaggatatcgagaacactctggctaagatggatgacgggcaactggacggcttggcattcggggccattcaactggacggggacggtaatattctccagtacaatgcagctgaaggtgacataactggacgtgatcctaaacaagtgattggcaagaatttcttcaaagatgttgctcccggcaccgattcacctgagttttacgggaagttcaaagagggcgttgcctccggcaacctcaacactatgttcgaatggatgataccgacaagcagaggcccaaccaaggtaaaaattcatatgaagaaagctctgtctggagactcttattgggtgtttgtgaagagagtcgaaaacctgtattttcagggcggctccggcggcgaacacgttgcctttggctctgaggatatcgaaaatacgctggccaaaatggatgacggtcagctggatggcctggcattcggagccatacagcttgacggcgatggaaacatcctccagtacaacgccgcggagggtgacattaccggtcgcgatccaaaacaggttattggcaagaacttctttaaggacgttgcacccgggactgactctcccgaattttacggaaagttcaaggaagggggtggcatctggcaatcttaatacaatgtttgaatggatgattcctactagtagaggaccaacaaaggtgaagatccacatgaagaaagccttgtccggcgactcctactgggtctttgtgaaacgggtgggatccgaacaaaagcttatttctgaagaggacttgtaa |
| pAG404 | HIS-$FAST_{25-125}$-ENLYFQGGSGG-$FAST_{1-24}$ | atgggcagcagccatcatcatcatcatcacagcagcggcctggtgccgcgcggcagccatatggctagcatgggcttggcattcggggccattcaactggacggggacggtaatattctccagtacaatgcagctgaaggtgacataactggacgtgatcctaaacaagtgattggcaagaatttcttcaaagatgttgctcccggcaccgattcacctgagttttacgggaagttcaaagagggcgttgcctccggcaacctcaacactatgttcgaatggatgataccgacaagcagaggcccaaccaaggtaaaaattcatatgaagaaagctctgtctggagactcttattgggtgtttgtgaagagagtcgaaaacctgtattttcagggcggctccggcggcgaacacgttgcctttggctctgaggatatcgaaaatacgctggccaaaatggatgacggtcagctggat |
| pAG405 | HIS-$FAST_{36-125}$-ENLYFQGGSGG-$FAST_{1-35}$ | atgggcagcagccatcatcatcatcatcacagcagcggcctggtgccgcgcggcagccatatggctagcatggacggtaatattctccagtacaatgcagctgaaggtgacataactggacgtgatcctaaacaagtgattggcaagaatttcttcaaagatgttgctcccggcaccgattcacctgagttttacgggaagttcaaagagggcgttgcctccggcaacctcaacactatgttcgaatggatgataccgacaagcagaggcccaaccaaggtaaaaattcatatgaagaaagctctgtctggagactcttattgggtgtttgtgaagagagtcgaaaacctgtattttcagggcggctccggcggcgaacacgttgcctttggctctgaggatatcgaaaatacgctggccaaaatggatgacggtcagctggatggcctggcattcggagccatacagcttgacggc |
| pAG406 | HIS-$FAST_{60-125}$-ENLYFQGGSGG-$FAST_{1-59}$ | atgggcagcagccatcatcatcatcatcacagcagcggcctggtgccgcgcggcagccatatggctagcatgaagaatttcttcaaagatgttgctcccggcaccgattcacctgagttttacgggaagttcaaagagggcgttgcctccggcaacctcaacactatgttcgaatggatgataccgacaagcagaggcccaaccaaggtaaaaattcatatgaagaaagctctgtctggagactcttattgggtgtttgtgaagagagtcgaaaacctgtattttcagggcggctccggcggcgaacacgttgcctttggctctgaggatatcgaaaatacgctggccaaaatggatgacggtcagctggatggcctggcattcggagccatacagcttgacggcgatggaaacatcctccagtacaacgccgcggagggtgacattaccggtcgcgatccaaaacaggttattggc |
| pAG407 | HIS-$FAST_{73-125}$-ENLYFQGGSGG-$FAST_{1-72}$ | atgggcagcagccatcatcatcatcatcacagcagcggcctggtgccgcgcggcagccatatggctagcatgcctgagttttacgggaagttcaaagagggcgttgcctccggcaacctcaacactatgttcgaatggatgataccgacaagcagaggcccaaccaaggtaaaaattcatatgaagaaagctctgtctggagactcttattgggtgtttgtgaagagagtcgaaaacctgtattttcagggcggctccggcggcgaacacgttgcctttggctctgaggatatcgaaaatacgctggccaaaatggatgacggtcagctggatggcctggcattcggagccatacagcttgacggcgatggaaacatcctccagtacaacgccgcggagggtgacattaccggtcgcgatccaaaacaggttattggcaagaacttctttaaggacgttgcacccgggactgactct |
| pAG408 | HIS-$FAST_{90-125}$-ENLYFQGGSGG-$FAST_{1-89}$ | atgggcagcagccatcatcatcatcatcacagcagcggcctggtgccgcgcggcagccatatggctagcatgactatgttcgaatggatgataccgacaagcagaggcccaaccaaggtaaaaattcatatgaagaaagctctgtctggagactcttattgggtgtttgtgaagagagtcgaaaacctgtattttcagggcggctccggcggcgaacacgttgcctttggctctgaggatatcgaaaatacgctggccaaaatggatgacggtcagctggatggcctggcattcggagccatacagcttgacggcgatggaaacatcctccagtacaacgccgcggagggtgacattaccggtcgcgatccaaaacaggttattggcaagaacttctttaaggacgttgcacccgggactgactctcccgaattttacggaaagttcaaggaagggggtggcatctggcaatcttaat |
| pAG409 | HIS-$FAST_{99-125}$-ENLYFQGGSGG-$FAST_{1-98}$ | atgggcagcagccatcatcatcatcatcacagcagcggcctggtgccgcgcggcagccatatggctagcatgagcagaggcccaaccaaggtaaaaattcatatgaagaaagctctgtctggagactcttattgggtgtttgtgaagagagtcgaaaacctgtattttcagggcggctccggcggcgaacacgttgcctttggctctgaggatatcgaaaatacgctggccaaaatggatgacggtcagctggatggcctggcattcggagccatacagcttgacggcgatggaaacatcctccagtacaacgccgcggagggtgacattaccggtcgcgatccaaaacaggttattggcaagaacttctttaaggacgttgcacccgggactgactctcccgaattttacggaaagttcaaggaagggggtggcatctggcaatcttaatacaatgtttgaatggatgattcctact |
| pAG410 | HIS-$FAST_{103-125}$-ENLYFQGGSGG-$FAST_{1-102}$ | atgggcagcagccatcatcatcatcatcacagcagcggcctggtgccgcgcggcagccatatggctagcatgaccaaggtaaaaattcatatgaagaaagctctgtctggagactcttattgggtgtttgtgaagagagtcgaaaacctgtattttcagggcggctccggcggcgaacacgttgcctttggctctgaggatatcgaaaatacgctggccaaaatggatgacggtcagctggatggcctggcattcggagccatacagcttgacggcgatggaaacatcctccagtacaacgccgcggagggtgacattaccggtcgcgatccaaaacaggttattggcaagaacttctttaaggacgttgcacccgggactgactctcccgaattttacggaaagttcaaggaagggggtggcatctggcaatcttaatacaatgtttgaatggatgattcctactagtagaggacca |
| pAG473 | HIS-$FAST_{25-125}$-GGSGGSGGSGG-$FAST_{1-24}$ | atgggcagcagccatcatcatcatcatcacagcagcggcctggtgccgcgcggcagccatatggctagcatgggcttggcattcggggccattcaactggacggggacggtaatattctccagtacaatgcagctgaaggtgacataactggacgtgatcctaaacaagtgattggcaagaatttcttcaaagatgttgctcccggcaccgattcacctgagttttacgggaagttcaaagagggcgttgcctccggcaacctcaacactatgttcgaatggatgataccgacaagcagaggcccaaccaaggtaaaaattcatatgaagaaagctctgtctggagactcttattgggtgtttgtgaagagagtcggcggctccggcggctccggcggctccggcggcgaacacgttgcctttggctctgaggatatcgaaaatacgctggccaaaatggatgacggtcagctggat |
| pAG475 | HIS-$FAST_{25-125}$-GGSGGSGG-$FAST_{1-24}$ | atgggcagcagccatcatcatcatcatcacagcagcggcctggtgccgcgcggcagccatatggctagcatgggcttggcattcggggccattcaactggacggggacggtaatattctccagtacaatgcagctgaaggtgacataactggacgtgatcctaaacaagtgattggcaagaatttcttcaaagatgttgctcccggcaccgattcacctgagttttacgggaagttcaaagagggcgttgcctccggcaacctcaacactatgttcgaatggatgataccgacaagcagaggcccaaccaaggtaaaaattcatatgaagaaagctctgtctggagactcttattgggtgtttgtgaagagagtcggcggctccggcggctccggcggcgaacacgttgcctttggctctgaggatatcgaaaatacgctggccaaaatggatgacggtcagctggat |
| pAG478 | HIS-$FAST_{25-125}$-GGSGG-$FAST_{1-24}$ | atgggcagcagccatcatcatcatcatcacagcagcggcctggtgccgcgcggcagccatatggctagcatgggcttggcattcggggccattcaactggacggggacggtaatattctccagtacaatgcagctgaaggtgacataactggacgtgatcctaaacaagtgattggcaagaatttcttcaaagatgttgctcccggcaccgattcacctgagttttacgggaagttcaaagagggcgttgcctccggcaacctcaacactatgttcgaatggatgataccgacaagcagaggcccaaccaaggtaaaaattcatatgaagaaagctctgtctggagactcttattgggtgtttgtgaagagagtcggcggctccggcggcgaacacgttgcctttggctctgaggatatcgaaaatacgctggccaaaatggatgacggtcagctggat |
| pAG480 | HIS-$FAST_{25-125}$-GGS-$FAST_{1-24}$ | atgggcagcagccatcatcatcatcatcacagcagcggcctggtgccgcgcggcagccatatggctagcatgggcttggcattcggggccattcaactggacggggacggtaatattctccagtacaatgcagctgaaggtgacataactggacgtgatcctaaacaagtgattggcaagaatttcttcaaagatgttgctcccggcaccgattcacctgagttttacgggaagttcaaagagggcgttgcctccggcaacctcaacactatgttcgaatggatgataccgacaagcagaggcccaaccaaggtaaaaattcatatgaagaaagctctgtctggagactcttattgggtgtttgtgaagagagtcggcggctccgaacacgttgcctttggctctgaggatatcgaaaatacgctggccaaaatggatgacggtcagctggat |
| pAG482 | HIS-$FAST_{25-125}$-$FAST_{1-24}$ | atgggcagcagccatcatcatcatcatcacagcagcggcctggtgccgcgcggcagccatatggctagcatgggcttggcattcggggccattcaactggacggggacggtaatattctccagtacaatgcagctgaaggtgacataactggacgtgatcctaaacaagtgattggcaagaatttcttcaaagatgttgctcccggcaccgattcacctgagttttacgggaagttcaaagagggcgttgcctccggcaacctcaacactatgttcgaatggatgataccgacaagcagaggcccaaccaaggtaaaaattcatatgaagaaagctctgtctggagactcttattgggtgtttgtgaagagagtcgaacacgttgcctttggctctgaggatatcgaaaatacgctggccaaaatggatgacggtcagctggat |
| pAG474 | HIS-$FAST_{99-125}$-GGSGGSGGSGG -$FAST_{1-98}$ | atgggcagcagccatcatcatcatcatcacagcagcggcctggtgccgcgcggcagccatatggctagcatgAGCAGAGGCCCAACCAAGGTAAAAaTTCATATGAAGAAAGCTCTGTCTGGAGACTCTTATTGGGTGTTTGTGAAGAGAGTCggcggctccggcggctccggcggctccggcggcGAACACGTTGCCTTTGGCTCTGAGGATATCGAAAATACGCTGGCCAAAATGGATGACGGTCAGCTGGATGGCCTGGCATTCGGAGCCATACAGCTTGACGGCGATGGAAACATCCTCCAGTACAACGCCGCGGAGGGTGACATTACCGGTCGCGATCCAAAACAGGTTATTGGCAAGAACTTCTTTAAGGACGTTGCACCCGGGACTGACTCTCCCGAATTTTACGGAAAGTTCAAGGAAGGGGTGGCATCTGGCAATCTTAATACAATGTTTGAATGGATGATTCCTACT |
| pAG476 | HIS-$FAST_{99-125}$-GGSGGSGG -$FAST_{1-98}$ | atgggcagcagccatcatcatcatcatcacagcagcggcctggtgccgcgcggcagccatatggctagcatgAGCAGAGGCCCAACCAAGGTAAAAaTTCATATGAAGAAAGCTCTGTCTGGAGACTCTTATTGGGTGTTTGTGAAGAGAGTCggcggctccggcggctccggcggcGAACACGTTGCCTTTGGCTCTGAGGATATCGAAAATACGCTGGCCAAAATGGATGACGGTCAGCTGGATGGCCTGGCATTCGGAGCCATACAGCTTGACGGCGATGGAAACATCCTCCAGTACAACGCCGCGGAGGGTGACATTACCGGTCGCGATCCAAAACAGGTTATTGGCAAGAACTTCTTTAAGGACGTTGCACCCGGGACTGACTCTCCCGAATTTTACGGAAAGTTCAAGGAAGGGGTGGCATCTGGCAATCTTAATACAATGTTTGAATGGATGATTCCTACT |
| pAG479 | HIS-$FAST_{99-125}$-GGSGG-$FAST_{1-98}$ | atgggcagcagccatcatcatcatcatcacagcagcggcctggtgccgcgcggcagccatatggctagcatgAGCAGAGGCCCAACCAAGGTAAAAaTTCATATGAAGAAAGCTCTGTCTGGAGACTCTTATTGGGTGTTTGTGAAGAGAGTCggcggctccggcggcGAACACGTTGCCTTTGGCTCTGAGGATATCGAAAATACGCTGGCCAAAATGGATGACGGTCAGCTGGATGGCCTGGCATTCGGAGCCATACAGCTTGACGGCGATGGAAACATCCTCCAGTACAACGCCGCGGAGGGTGACATTACCGGTCGCGATCCAAAACAGGTTATTGGCAAGAACTTCTTTAAGGACGTTGCACCCGGGACTGACTCTCCCGAATTTTACGGAAAGTTCAAGGAAGGGGTGGCATCTGGCAATCTTAATACAATGTTTGAATGGATGATTCCTACT |

| | | |
|---|---|---|
| pAG481 | HIS-FAST$_{99-125}$-GGS-FAST$_{1-98}$ | atgggcagcagccatcatcatcatcatcacagcagcggcctggtgccgcgcggcagccatatggctagcatgAGCAGAGGCCCAACCAAGGTA<br>AAAaTTCATATGAAGAAAGCTCTGTCTGGAGACTCTTATTGGGTGTTTGTGAAGAGAGTCggcggctccGAACACGTTGCCTTTGGCTCTGAG<br>GATATCGAAAATACGCTGGCCAAAATGGATGACGGTCAGCTGGATGGCCTGGCATTCGGAGCCATACAGCTTGACGGCGATGGAAACATCCTC<br>CAGTACAACGCCGCGGAGGGTGACATTACCGGTCGCGATCCAAAACAGGTTATTGGCAAGAACTTCTTTAAGGACGTTGCACCCGGGACTGAC<br>TCTCCCGAATTTTACGGAAAGTTCAAGGAAGGGGTGGCATCTGGCAATCTTAATACAATGTTTGAATGGATGATTCCTACT |
| pAG483 | HIS-FAST$_{99-125}$-FAST$_{1-98}$ | atgggcagcagccatcatcatcatcatcacagcagcggcctggtgccgcgcggcagccatatggctagcatgAGCAGAGGCCCAACCAAGGTA<br>AAAaTTCATATGAAGAAAGCTCTGTCTGGAGACTCTTATTGGGTGTTTGTGAAGAGAGTCGAACACGTTGCCTTTGGCTCTGAGGATATCGAA<br>AATACGCTGGCCAAAATGGATGACGGTCAGCTGGATGGCCTGGCATTCGGAGCCATACAGCTTGACGGCGATGGAAACATCCTCCAGTACAAC<br>GCCGCGGAGGGTGACATTACCGGTCGCGATCCAAAACAGGTTATTGGCAAGAACTTCTTTAAGGACGTTGCACCCGGGACTGACTCTCCCGAA<br>TTTTACGGAAAGTTCAAGGAAGGGGTGGCATCTGGCAATCTTAATACAATGTTTGAATGGATGATTCCTACT |
| pAG978 | HIS-nanoFAST | atgggcagcagccatcatcatcatcatcacagcagcggcctggtgccgcgcggcagccatatggctagcgaaaacctgtattttcagggcATG<br>TTTGGCGCAATTCAGCTCGATGGTGACGGAATATCCTGCAGTACAATGCTGCTGAAGGAGACATCACAGGCAGAGATCCCAAACAGGTGATT<br>GGGAAGAACTTCTTCAAGGATGTTGCACCTGGAACGGATTCTCCCGAGTTTTACGGCAAATTCAAGGAAGGCGTAGCGTCAGGGAATCTGAAC<br>ACCATGTTCGAATGGATGATACCGACAAGCAGGGGACCAACCAAGGTCAAGGTGCACATGAAGAAAGCCCTTTCCGGTGACAGCTATTGGGTC<br>TTTGTGAAACGGGTGTAA |
| pAG802 | MYC-FKBP-FAST$_{99-125}$-IRES-HA-iRFP670 | atggaacaaaagcttatttctgaagaggacttggaattcggagtgcaggtggaaaccatctccccaggagacgggcgcaccttccccaagcgc<br>ggccagacctgcgtggtgcactacaccgggatgcttgaagatggaaagaaatttgattcctcccgggacagaaacaagccctttaagtttatg<br>ctaggcaagcaggaggtgatccgaggctgggaagaaggggttgcccagatgagtgtgggtcagagagccaaactgactatatctccagattat<br>gcctatggtgccactgggcacccaggcatcatcccaccacatgccactctcgtcttcgatgtggagcttctaaaactggaagaatccggagga<br>ggcggcagcggcggaggggatccagcaggggaccaaccaaggtcaaggtgcacatgaagaaagccctttccggtgacagctattgggtcttt<br>gtgaaacgggtgtaactcgaggactacaaggacgacgacgacaagcccgggatccgcccctctccctcccccccccctaacgttactggccga<br>agccgcttggaataaggccggtgtgcgtttgtctatatgttattttccaccatattgccgtcttttggcaatgtgagggcccggaaacctggc<br>cctgtcttcttgacgagcattcctaggggtctttcccctctcgccaaaggaatgcaaggtctgttgaatgtcgtgaaggaagcagttcctctg<br>gaagcttcttgaagacaaacaacgtctgtagcgaccctttgcaggcagcggaaccccccacctggcgacaggtgcctctgcggccaaaagcca<br>cgtgtataagatacacctgcaaaggcggcacaaccccagtgccacgttgtgagttggatagttgtggaaagagtcaaatggctctcctcaagc<br>gtattcaacaaggggctgaaggatgcccagaaggtaccccattgtatgggatctgatctggggcctcggtgcacatgctttacatgtgtttag<br>tcgaggttaaaaaacgtctaggccccccgaaccacggggacgtggttttcctttgaaaaacacgatgataatatggccacaaccatgcgatc<br>gtacccatacgatgttccagattacgctgaattcatggcgcgtaaggtcgatctcacctcctgcgatcgcgagccgatccacatccccggcag<br>cattcagccgtgcggctgcctgctagcctgcgacgcgcaggcggtgcggatcacgcgcattacggaaaatgccggcgcgttctttggacgcga<br>aactccgcgggtcggtgagctactcgccgattacttcggcgagaccgaagcccatgcgctgcgcaacgcactggcgcagtcttccgatccaaa<br>gcgaccggcgctgatcttcggttggcgcgacggcctgaccggccgcaccttcgacatctcactgcatcgccatgacggtacatcgatcatcga<br>gttcgagcctgcggcggccgaacaggccgacaatccgctgcggctgacgcggcagatcatcgcgcgcaccaaagaactgaagtcgctcgaaga<br>gatggccgcacgggtgccgcgctatctgcaggcgatgctcggctatcaccgcgtgatgttgtaccgcttcgcggacgacggctccgggatggt<br>gatcggcgaggcgaagcgcagcgacctcgagagctttctcggtcagcactttccggcgtcgctggtcccgcagcaggcgcggctactgtactt<br>gaagaacgcgatccgcgtggtctcggattcgcgcggcatcagcagccggatcgtgcccgagcacgacgcctccggcgccgcgctcgatctgtc<br>gttcgcgcacctgcgcagcatctcgccctgccatctcgaatttctgcggaacatgggcgtcagcgcctcgatgtcgctgtcgatcatcattga<br>cggcacgctatggggattgatcatctgtcatcattacgagccgcgtgccgtgccgatggcgcagcgcgtcgcggccgaaatgttcgccgactt<br>cttatcgctgcacttcaccgccgcccaccaccaacgc |
| pAG935 | FAST$_{1-98}$-FRB-MYC-IRES-HA-mTurquoise2 | atggagcatgttgcctttggcagtgaggacatcgagaacactctggccaaaatggacgacggacaactggatgggttggcctttggcgcaatt<br>cagctcgatggtgacggaatatcctgcagtacaatgctgctgaaggagacatcacaggcagagatcccaaacaggtgattgggaagaacttc<br>ttcaaggatgttgcacctggaacggattctcccgagttttacggcaaattcaaggaaggcgtagcgtcagggaatctgaacaccatgttcgaa<br>tggatgataccgacatccggaggaggcggcagcggcggaggggatccatgtggcatgaaggcctggaagaggcatctcgtttgtactttggg<br>gaaaggaacgtgaaaggcatgtttgaggtgctggagcccttgcatgctatgatggaacggggccccagactctgaaggaaacatcctttaat<br>caggcctatggtcgagatttaatggaggcccaagagtggtgcaggaagtacatgaaatcagggaatgtcaaggacctcacccaagcctgggac<br>ctctattatcatgtgttccgacgaatctcaaagcaggtcgaattcgaacaaaagcttatttctgaagaggacttgtaactcgaggactacaag<br>gacgacgacgacaagcccgggatccgcccctctccctcccccccccctaacgttactggccgaagccgcttggaataaggccggtgtgcgttt<br>gtctatatgttattttccaccatattgccgtcttttggcaatgtgagggcccggaaacctggccctgtcttcttgacgagcattcctaggggt<br>ctttcccctctcgccaaaggaatgcaaggtctgttgaatgtcgtgaaggaagcagttcctctggaagcttcttgaagacaaacaacgtctgta<br>gcgaccctttgcaggcagcggaaccccccacctggcgacaggtgcctctgcggccaaaagccacgtgtataagatacacctgcaaaggcggca<br>caaccccagtgccacgttgtgagttggatagttgtggaaagagtcaaatggctctcctcaagcgtattcaacaaggggctgaaggatgcccag<br>aaggtaccccattgtatgggatctgatctggggcctcggtgcacatgctttacatgtgtttagtcgaggttaaaaaacgtctaggccccccg<br>aaccacggggacgtggttttcctttgaaaaacacgatgataatatggccacaaccatgcgatcgtacccatacgatgttccagattacgctga<br>attcatggtgagcaagggcgaggagctgttcaccggggtggtgcccatcctggtcgagctggacggcgacgtaaacggccacaagttcagcgt<br>gtccggcgagggcgagggcgatgccacctacggcaagctgaccctgaagttcatctgcaccaccggcaagctgcccgtgccctggcccaccct<br>cgtgaccaccctgtcctggggcgtgcagtgcttcgcccgctaccccgaccacatgaagcagcacgacttcttcaagtccgccatgcccgaagg<br>ctacgtccaggagcgcaccatcttcttcaaggacgacggcaactacaagacccgcgccgaggtgaagttcgagggcgacaccctggtgaaccg<br>catcgagctgaagggcatcgacttcaaggaggacggcaacatcctggggcacaagctggagtacaactacttcagcgacaacgtctatatcac<br>cgccgacaagcagaagaacggcatcaaggccaacttcaagatccgccacaacatcgaggacggcggcgtgcagctcgccgaccactaccagca<br>gaacacccccatcggcgacggccccgtgctgctgcccgacaaccactacctgagcacccagtccaagctgagcaaagaccccaacgagaagcg<br>cgatcacatggtcctgctggagttcgtgaccgccgccgggatcactctcggcatggacgagctgtacaagtaa |
| pAG974 | MYC-FKBP-$^{Rspa}$FAST$_{99-125}$-IRES-HA-iRFP670 | atggaacaaaagcttatttctgaagaggacttggaattcggagtgcaggtggaaaccatctccccaggagacgggcgcaccttccccaagcgc<br>ggccagacctgcgtggtgcactacaccgggatgcttgaagatggaaagaaatttgattcctcccgggacagaaacaagccctttaagtttatg<br>ctaggcaagcaggaggtgatccgaggctgggaagaaggggttgcccagatgagtgtgggtcagagagccaaactgactatatctccagattat<br>gcctatggtgccactgggcacccaggcatcatcccaccacatgccactctcgtcttcgatgtggagcttctaaaactggaagaatccggagga<br>ggcggcagcggcggaggggatccagcagaggccccaccaaggtgaaggtgcacatgaagaaggccatgaccggcgacagcttctggatcttc<br>gtgaagagactgtaactcgaggactacaaggacgacgacgacaagcccgggatccgcccctctccctcccccccccctaacgttactggccga<br>agccgcttggaataaggccggtgtgcgtttgtctatatgttattttccaccatattgccgtcttttggcaatgtgagggcccggaaacctggc<br>cctgtcttcttgacgagcattcctaggggtctttcccctctcgccaaaggaatgcaaggtctgttgaatgtcgtgaaggaagcagttcctctg<br>gaagcttcttgaagacaaacaacgtctgtagcgaccctttgcaggcagcggaaccccccacctggcgacaggtgcctctgcggccaaaagcca<br>cgtgtataagatacacctgcaaaggcggcacaaccccagtgccacgttgtgagttggatagttgtggaaagagtcaaatggctctcctcaagc<br>gtattcaacaaggggctgaaggatgcccagaaggtaccccattgtatgggatctgatctggggcctcggtgcacatgctttacatgtgtttag<br>tcgaggttaaaaaacgtctaggccccccgaaccacggggacgtggttttcctttgaaaaacacgatgataatatggccacaaccatgcgatc<br>gtacccatacgatgttccagattacgctgaattcatggcgcgtaaggtcgatctcacctcctgcgatcgcgagccgatccacatccccggcag<br>cattcagccgtgcggctgcctgctagcctgcgacgcgcaggcggtgcggatcacgcgcattacggaaaatgccggcgcgttctttggacgcga<br>aactccgcgggtcggtgagctactcgccgattacttcggcgagaccgaagcccatgcgctgcgcaacgcactggcgcagtcttccgatccaaa<br>gcgaccggcgctgatcttcggttggcgcgacggcctgaccggccgcaccttcgacatctcactgcatcgccatgacggtacatcgatcatcga<br>gttcgagcctgcggcggccgaacaggccgacaatccgctgcggctgacgcggcagatcatcgcgcgcaccaaagaactgaagtcgctcgaaga<br>gatggccgcacgggtgccgcgctatctgcaggcgatgctcggctatcaccgcgtgatgttgtaccgcttcgcggacgacggctccgggatggt<br>gatcggcgaggcgaagcgcagcgacctcgagagctttctcggtcagcactttccggcgtcgctggtcccgcagcaggcgcggctactgtactt<br>gaagaacgcgatccgcgtggtctcggattcgcgcggcatcagcagccggatcgtgcccgagcacgacgcctccggcgccgcgctcgatctgtc<br>gttcgcgcacctgcgcagcatctcgccctgccatctcgaatttctgcggaacatgggcgtcagcgcctcgatgtcgctgtcgatcatcattga<br>cggcacgctatggggattgatcatctgtcatcattacgagccgcgtgccgtgccgatggcgcagcgcgtcgcggccgaaatgttcgccgactt<br>cttatcgctgcacttcaccgccgcccaccaccaacgc |
| pAG1095 | $^{Rspa}$FAST$_{1-98}$-FRB-MYC-IRES-HA-mTurquoise2 | atggagaccgtgagattcggcggcgacgacatcgagaacagcctggccaagatggacgacaaggccctggacaagctggccttcggcgccatc<br>cagctggacggcaacggcaagatcatccactacaacgccgccgagggcaccatcaccggcagagaccccaagaccgtgatcggcaagaacttc<br>ttcaccgacgtggcccccggcacccagagcaaggagttccagggcagattcaaggagggcgtgcagaagggcgacctgaacaccatgttcgag<br>tggatgatccccacctccggaggaggcggcagcggcggaggggatccatgtggcatgaaggcctggaagaggcatctcgtttgtactttggg<br>gaaaggaacgtgaaaggcatgtttgaggtgctggagcccttgcatgctatgatggaacggggccccagactctgaaggaaacatcctttaat<br>caggcctatggtcgagatttaatggaggcccaagagtggtgcaggaagtacatgaaatcagggaatgtcaaggacctcacccaagcctgggac<br>ctctattatcatgtgttccgacgaatctcaaagcaggtcgaattcgaacaaaagcttatttctgaagaggacttgtaactcgaggactacaag<br>gacgacgacgacaagcccgggatccgcccctctccctcccccccccctaacgttactggccgaagccgcttggaataaggccggtgtgcgttt<br>gtctatatgttattttccaccatattgccgtcttttggcaatgtgagggcccggaaacctggccctgtcttcttgacgagcattcctaggggt<br>ctttcccctctcgccaaaggaatgcaaggtctgttgaatgtcgtgaaggaagcagttcctctggaagcttcttgaagacaaacaacgtctgta<br>gcgaccctttgcaggcagcggaaccccccacctggcgacaggtgcctctgcggccaaaagccacgtgtataagatacacctgcaaaggcggca<br>caaccccagtgccacgttgtgagttggatagttgtggaaagagtcaaatggctctcctcaagcgtattcaacaaggggctgaaggatgcccag<br>aaggtaccccattgtatgggatctgatctggggcctcggtgcacatgctttacatgtgtttagtcgaggttaaaaaacgtctaggccccccg<br>aaccacggggacgtggttttcctttgaaaaacacgatgataatatggccacaaccatgcgatcgtacccatacgatgttccagattacgctga<br>attcatggtgagcaagggcgaggagctgttcaccggggtggtgcccatcctggtcgagctggacggcgacgtaaacggccacaagttcagcgt<br>gtccggcgagggcgagggcgatgccacctacggcaagctgaccctgaagttcatctgcaccaccggcaagctgcccgtgccctggcccaccct<br>cgtgaccaccctgtcctggggcgtgcagtgcttcgcccgctaccccgaccacatgaagcagcacgacttcttcaagtccgccatgcccgaagg<br>ctacgtccaggagcgcaccatcttcttcaaggacgacggcaactacaagacccgcgccgaggtgaagttcgagggcgacaccctggtgaaccg<br>catcgagctgaagggcatcgacttcaaggaggacggcaacatcctggggcacaagctggagtacaactacttcagcgacaacgtctatatcac |

| | | |
|---|---|---|
| | | cgccgacaagcagaagaacggcatcaaggccaacttcaagatccgccacaacatcgaggacggcggcgtgcagctcgccgaccactaccagca<br>gaacacccccatcggcgacggccccgtgctgctgcccgacaaccactacctgagcacccagtccaagctgagcaaagaccccaacgagaagcg<br>cgatcacatggtcctgctggagttcgtgaccgccgccgggatcactctcggcatggacgagctgtacaagtaa |

| Plasmid | ORF | ORF sequence |
|---|---|---|
| pAG1127 | MYC-$^{RspA}$FAST$_{99-114}$-FRB-$^{RspA}$FAST$_{1-98}$-IRES-HA-mTurquoise2 | atggaacaaaagcttatttctgaagaggacttggaattcagcagaggccccaccaaggtgaaggtgcacatgaagaaggccatgaccagcggc<br>gggggaggctccggaggcggaggcagcatgtggcatgaaggcctggaagaggcatctcgtttgtactttggggaaaggaacgtgaaaggcatg<br>tttgaggtgctggagcccttgcatgctatgatggaacggggcccccagactctgaaggaaacatcctttaatcaggcctatggtcgagattta<br>atggaggcccaagagtggtgcaggaagtacatgaaatcagggaatgtcaaggacctcacccaagcctgggacctctattatcatgtgttccga<br>cgaatctcaaagcaggtctccggaggaggcggcagcggcggagggggatccatggagaccgtgagattcggcggcgacgacatcgagaacagc<br>ctggccaagatggacgacaaggccctggacaagctggccttcggcgccatccagctggacggcaacggcaagatcatccactacaacgccgcc<br>gagggcaccatcaccggcagagaccccaagaccgtgatcggcaagaacttcttcaccgacgtggccccggcacccagagcaaggagttccag<br>ggcagattcaaggaggcgtgcagaagggcgacctgaacaccatgttcgagtggatgatccccacctaactcgaggactacaaggacgacgac<br>gacaagcccgggatccgcccctctccctcccccccccctaacgttactggccgaagccgcttggaataaggccggtgtgcgtttgtctatatg<br>ttattttccaccatattgccgtcttttggcaatgtgagggcccggaaacctggccctgtcttcttgacgagcattcctaggggtctttcccct<br>ctcgccaaaggaatgcaaggtctgttgaatgtcgtgaaggaagcagttcctctggaagcttcttgaagacaaacaacgtctgtagcgaccctt<br>tgcaggcagcggaaccccccacctggcgacaggtgcctctgcggccaaaagccacgtgtataagatacacctgcaaaggcggcacaaccccag<br>tgccacgttgtgagttggatagttgtggaaagagtcaaatggctctcctcaagcgtattcaacaaggggctgaaggatgcccagaaggtaccc<br>cattgtatgggatctgatctggggcctcggtgcacatgctttacatgtgtttagtcgaggttaaaaaaacgtctaggccccccgaaccacggg<br>gacgtggttttcctttgaaaaacacgatgataatatggccacaaccatgcgatcgtacccatacgatgttccagattacgctgaattcatggt<br>gagcaagggcgaggagctgttcaccggggtggtgcccatcctggtcgagctggacggcgacgtaaacggccacaagttcagcgtgtccggcga<br>gggcgagggcgatgccacctacggcaagctgaccctgaagttcatctgcaccaccggcaagctgcccgtgccctggcccaccctcgtgaccac<br>cctgtcctggggcgtgcagtgcttcgcccgctaccccgaccacatgaagcagcacgacttcttcaagtccgccatgcccgaaggctacgtcca<br>ggagcgcaccatcttcttcaaggacgacggcaactacaagacccgcgccgaggtgaagttcgagggcgacaccctggtgaaccgcatcgagct<br>gaagggcatcgacttcaaggaggacggcaacatcctggggcacaagctggagtacaactacttcagcgacaacgtctatatcaccgccgacaa<br>gcagaagaacggcatcaaggccaacttcaagatccgccacaacatcgaggacggcggcgtgcagctcgccgaccactaccagcagaacacccc<br>catcggcgacggccccgtgctgctgcccgacaaccactacctgagcacccagtccaagctgagcaaagaccccaacgagaagcgcgatcacat<br>ggtcctgctggagttcgtgaccgccgccgggatcactctcggcatggacgagctgtacaagtaa |
| pAG580 | MYC-FKBP-$^{RspA}$FAST$_{115-125}$-IRES-HA-IRFP670 | atggaacaaaagcttatttctgaagaggacttggaattcggagtgcaggtggaaaccatctccccaggagacgggcgcaccttccccaagcgc<br>ggccagacctgcgtggtgcactacaccgggatgcttgaagatggaaagaaatttgattcctcccgggacagaaacaagccctttaagtttatg<br>ctaggcaagcaggaggtgatccgaggctgggaagaaggggttgcccagatgagtgtgggtcagagagccaaactgactatatctccagattat<br>gcctatggtgccactgggcacccaggcatcatcccaccacatgccactctcgtcttcgatgtggagcttctaaaactggaagaatccggagga<br>ggcggcagcggcggagggggatccggcgacagcttctggatcttcgtgaagagactgtaactcgaggactacaaggacgacgacgacaagccc<br>gggatccgcccctctccctcccccccccctaacgttactggccgaagccgcttggaataaggccggtgtgcgtttgtctatatgttattttcc<br>accatattgccgtcttttggcaatgtgagggcccggaaacctggccctgtcttcttgacgagcattcctaggggtctttcccctctcgccaaa<br>ggaatgcaaggtctgttgaatgtcgtgaaggaagcagttcctctggaagcttcttgaagacaaacaacgtctgtagcgaccctttgcaggcag<br>cggaaccccccacctggcgacaggtgcctctgcggccaaaagccacgtgtataagatacacctgcaaaggcggcacaaccccagtgccacgtt<br>gtgagttggatagttgtggaaagagtcaaatggctctcctcaagcgtattcaacaaggggctgaaggatgcccagaaggtaccccattgtatg<br>ggatctgatctggggcctcggtgcacatgctttacatgtgtttagtcgaggttaaaaaaacgtctaggccccccgaaccacggggacgtggtt<br>ttcctttgaaaaacacgatgataatatggccacaaccatgcgatcgtacccatacgatgttccagattacgctgaattcatggcgcgtaaggt<br>cgatctcacctcctgcgatcgcgagccgatccacatccccggcagcattcagccgtgcggctgcctgctagcctgcgacgcgcaggcggtgcg<br>gatcacgcgcattacggaaaatgccggcgcgttctttggacgcgaaactccgcgggtcggtgagctactcgccgattacttcggcgagaccga<br>agcccatgcgctgcgcaacgcactggcgcagtcttccgatccaaagcgaccggcgctgatcttcggttggcgcgacggcctgaccggccgcac<br>cttcgacatctcactgcatcgccatgacggtacatcgatcatcgagttcgagcctgcggcggccgaacaggccgacaatccgctgcggctgac<br>gcggcagatcatcgcgcgcaccaaagaactgaagtcgctcgaagagatggccgcacgggtgccgcgctatctgcaggcgatgctcggctatca<br>ccgcgtgatgttgtaccgcttcgcggacgacggctccgggatggtgatcggcgaggcgaagcgcagcgacctcgagagctttctcggtcagca<br>ctttccggcgtcgctggtcccgcagcaggcgcggctactgtacttgaagaacgcgatccgcgtggtctcggattcgcgcggcatcagcagccg<br>gatcgtgcccgagcacgacgcctccggcgccgcgctcgatctgtcgttcgcgcacctgcgcagcatctcgccctgccatctcgaatttctgcg<br>gaacatgggcgtcagcgcctcgatgtcgctgtcgatcatcattgacggcacgctatggggattgatcatctgtcatcattacgagccgcgtgc<br>cgtgccgatggcgcagcgcgtcgcggccgaaatgttcgccgacttcttatcgctgcacttcaccgccgcccaccaccaacgcgaacaaaagct<br>tatttctgaagaggacttgtaa |
| pAG1148 | MYC-pFAST$_{99-114}$-FRB-pFAST$_{1-98}$-IRES-HA-mTurquoise2 | atggaacaaaagcttatttctgaagaggacttggaattcagcaggggaccaaccaaggtcaaggtgcacttgaagaaagccctttccagcggc<br>gggggaggctccggaggcggaggcagcgagatgtggcatgaaggcctggaagaggcatctcgtttgtactttggggaaaggaacgtgaaaggc<br>atgtttgaggtgctggagcccttgcatgctatgatggaacggggcccccagactctgaaggaaacatcctttaatcaggcctatggtcgagat<br>ttaatggaggcccaagagtggtgcaggaagtacatgaaatcagggaatgtcaaggacctcacccaagcctgggacctctattatcatgtgttc<br>cgacgaatctcaaagcaggtctccggaggaggcggcagcggcggagggggatccatggagcatgttgcctttggcagtgaggacatcgagaac<br>actctggccaatatggacgacgaacaactggataggttggcctttggcgtaattcagctcgatggtgacgggaatatcctgctgtacaatgct<br>gctgaaggggacatcactggcagagatcccaaacaggtgattggaagaacttcttcaaggatgttgcacctggaacggatactcccgagttt<br>tacggcaaattcaaggaaggcgcagcgtcagggaatctgaacaccatgttcgaatggacgataccgacataactcgaggactacaaggacgac<br>gacgacaagcccgggatccgcccctctccctcccccccccctaacgttactggccgaagccgcttggaataaggccggtgtgcgtttgtctat<br>atgttattttccaccatattgccgtcttttggcaatgtgagggcccggaaacctggccctgtcttcttgacgagcattcctaggggtctttcc<br>cctctcgccaaaggaatgcaaggtctgttgaatgtcgtgaaggaagcagttcctctggaagcttcttgaagacaaacaacgtctgtagcgacc<br>ctttgcaggcagcggaaccccccacctggcgacaggtgcctctgcggccaaaagccacgtgtataagatacacctgcaaaggcggcacaaccc<br>cagtgccacgttgtgagttggatagttgtggaaagagtcaaatggctctcctcaagcgtattcaacaaggggctgaaggatgcccagaaggta<br>ccccattgtatgggatctgatctggggcctcggtgcacatgctttacatgtgtttagtcgaggttaaaaaaacgtctaggccccccgaaccac<br>ggggacgtggttttcctttgaaaaacacgatgataatatggccacaaccatgcgatcgtacccatacgatgttccagattacgctgaattcat<br>ggtgagcaagggcgaggagctgttcaccggggtggtgcccatcctggtcgagctggacggcgacgtaaacggccacaagttcagcgtgtccgg<br>cgagggcgagggcgatgccacctacggcaagctgaccctgaagttcatctgcaccaccggcaagctgcccgtgccctggcccaccctcgtgac<br>caccctgtcctggggcgtgcagtgcttcgcccgctaccccgaccacatgaagcagcacgacttcttcaagtccgccatgcccgaaggctacgt<br>ccaggagcgcaccatcttcttcaaggacgacggcaactacaagacccgcgccgaggtgaagttcgagggcgacaccctggtgaaccgcatcga<br>gctgaagggcatcgacttcaaggaggacggcaacatcctggggcacaagctggagtacaactacttcagcgacaacgtctatatcaccgccga<br>caagcagaagaacggcatcaaggccaacttcaagatccgccacaacatcgaggacggcggcgtgcagctcgccgaccactaccagcagaacac<br>ccccatcggcgacggccccgtgctgctgcccgacaaccactacctgagcacccagtccaagctgagcaaagaccccaacgagaagcgcgatca<br>catggtcctgctggagttcgtgaccgccgccgggatcactctcggcatggacgagctgtacaagtaa |
| pAG1151 | MYC-FKBP-pFAST$_{115-125}$-IRES-HA-IRFP670 | atggaacaaaagcttatttctgaagaggacttggaattcggagtgcaggtggaaaccatctccccaggagacgggcgcaccttccccaagcgc<br>ggccagacctgcgtggtgcactacaccgggatgcttgaagatggaaagaaatttgattcctcccgggacagaaacaagccctttaagtttatg<br>ctaggcaagcaggaggtgatccgaggctgggaagaaggggttgcccagatgagtgtgggtcagagagccaaactgactatatctccagattat<br>gcctatggtgccactgggcacccaggcatcatcccaccacatgccactctcgtcttcgatgtggagcttctaaaactggaagaatccggagga<br>ggcggcagcggcggagggggatccggtgacagatattgggtctttgtgaaacgggtgtaactcgaggactacaaggacgacgacgacaagccc<br>gggatccgcccctctccctcccccccccctaacgttactggccgaagccgcttggaataaggccggtgtgcgtttgtctatatgttattttcc<br>accatattgccgtcttttggcaatgtgagggcccggaaacctggccctgtcttcttgacgagcattcctaggggtctttcccctctcgccaaa<br>ggaatgcaaggtctgttgaatgtcgtgaaggaagcagttcctctggaagcttcttgaagacaaacaacgtctgtagcgaccctttgcaggcag<br>cggaaccccccacctggcgacaggtgcctctgcggccaaaagccacgtgtataagatacacctgcaaaggcggcacaaccccagtgccacgtt<br>gtgagttggatagttgtggaaagagtcaaatggctctcctcaagcgtattcaacaaggggctgaaggatgcccagaaggtaccccattgtatg<br>ggatctgatctggggcctcggtgcacatgctttacatgtgtttagtcgaggttaaaaaaacgtctaggccccccgaaccacggggacgtggtt<br>ttcctttgaaaaacacgatgataatatggccacaaccatgcgatcgtacccatacgatgttccagattacgctgaattcatggcgcgtaaggt<br>cgatctcacctcctgcgatcgcgagccgatccacatccccggcagcattcagccgtgcggctgcctgctagcctgcgacgcgcaggcggtgcg<br>gatcacgcgcattacggaaaatgccggcgcgttctttggacgcgaaactccgcgggtcggtgagctactcgccgattacttcggcgagaccga<br>agcccatgcgctgcgcaacgcactggcgcagtcttccgatccaaagcgaccggcgctgatcttcggttggcgcgacggcctgaccggccgcac<br>cttcgacatctcactgcatcgccatgacggtacatcgatcatcgagttcgagcctgcggcggccgaacaggccgacaatccgctgcggctgac<br>gcggcagatcatcgcgcgcaccaaagaactgaagtcgctcgaagagatggccgcacgggtgccgcgctatctgcaggcgatgctcggctatca<br>ccgcgtgatgttgtaccgcttcgcggacgacggctccgggatggtgatcggcgaggcgaagcgcagcgacctcgagagctttctcggtcagca<br>ctttccggcgtcgctggtcccgcagcaggcgcggctactgtacttgaagaacgcgatccgcgtggtctcggattcgcgcggcatcagcagccg<br>gatcgtgcccgagcacgacgcctccggcgccgcgctcgatctgtcgttcgcgcacctgcgcagcatctcgccctgccatctcgaatttctgcg |

| | | |
|---|---|---|
| | | gaacatgggcgtcagcgcctcgatgtcgctgtcgatcatcattgacggcacgctatggggattgatcatctgtcatcattacgagccgcgtgc<br>cgtgccgatggcgcagcgcgtcgcggccgaaatgttcgccgacttcttatcgctgcacttcaccgccgcccaccaccaacgctaa |
| pAG1184 | MYC-pFAST(99-114)-$^{N}$FRB-P2A-$^{C}$FRB-pFAST$_{1-98}$-IRES-HA-mTurquoise2 | atggaacaaaagcttatttctgaagaggacttggaattcagcaggggaccaaccaaggtcaaggtgcacttgaagaaagccctttccagcggc<br>gggggaggctccggaggcggaggcagcgagatgtggcatgaaggcctggaagaggcatctcgtttgtactttggggaaaggaacgtgaaaggc<br>atgtttgaggtgctggagcccttgcatgctatgatggaacggggccccggaagcggagctactaacttcagcctgctgaagcaggctggagac<br>gtggaggagaaccctggacctcagactctgaaggaaacatcctttaatcaggcctatggtcgagatttaatggaggcccaagagtggtgcagg<br>aagtacatgaaatcagggaatgtcaaggacctcacccaagcctgggacctctattatcatgtgttccgacgaatctcaaagcaggtctccgga<br>ggaggcggcagcggcggagggggatccatggagcatgttgcctttggcagtgaggacatcgagaacactctggccaatatggacgacgaacaa<br>ctggataggttggcctttggcgtaattcagctcgatggtgacgggaatatcctgctgtacaatgctgctgaaggggacatcactggcagagat<br>cccaaacaggtgattgggaagaacttcttcaaggatgttgcacctggaacggatactcccgagttttacggcaaattcaaggaaggcgcagcg<br>tcagggaatctgaacaccatgttcgaatggacgataccgacataactcgaggactacaaggacgacgacgacaagcccgggatccgcccctct<br>ccctcccccccccctaacgttactggccgaagccgcttggaataaggccggtgtgcgtttgtctatatgttattttccaccatattgccgtct<br>tttggcaatgtgagggcccggaaacctggccctgtcttcttgacgagcattcctaggggtctttcccctctcgccaaaggaatgcaaggtctg<br>ttgaatgtcgtgaaggaagcagttcctctggaagcttcttgaagacaaacaacgtctgtagcgaccctttgcaggcagcggaaccccccacct<br>ggcgacaggtgcctctgcggccaaaagccacgtgtataagatacacctgcaaaggcggcacaaccccagtgccacgttgtgagttggatagtt<br>gtggaaagagtcaaatggctctcctcaagcgtattcaacaaggggctgaaggatgcccagaaggtaccccattgtatgggatctgatctgggg<br>cctcggtgcacatgctttacatgtgtttagtcgaggttaaaaaacgtctaggccccccgaaccacggggacgtggttttcctttgaaaaaca<br>cgatgataatatggccacaaccatgcgatcgtacccatacgatgttccagattacgctgaattcatggtgagcaagggcgaggagctgttcac<br>cggggtggtgcccatcctggtcgagctggacggcgacgtaaacggccacaagttcagcgtgtccggcgagggcgagggcgatgccacctacgg<br>caagctgaccctgaagttcatctgcaccaccggcaagctgcccgtgccctggcccaccctcgtgaccaccctgtcctggggcgtgcagtgctt<br>cgcccgctaccccgaccacatgaagcagcacgacttcttcaagtccgccatgcccgaaggctacgtccaggagcgcaccatcttcttcaagga<br>cgacggcaactacaagacccgcgccgaggtgaagttcgagggcgacaccctggtgaaccgcatcgagctgaagggcatcgacttcaaggagga<br>cggcaacatcctggggcacaagctggagtacaactacttcagcgacaacgtctatatcaccgccgacaagcagaagaacggcatcaaggccaa<br>cttcaagatccgccacaacatcgaggacggcggcgtgcagctcgccgaccactaccagcagaacacccccatcggcgacggccccgtgctgct<br>gcccgacaaccactacctgagcacccagtccaagctgagcaaagaccccaacgagaagcgcgatcacatggtcctgctggagttcgtgaccgc<br>cgccgggatcactctcggcatggacgagctgtacaagtaa |
| pAG1223 | Lyn11-mCherry-$^{C}$FRB-pFAST$_{1-98}$ | atgggctgcatcaagtccaagggcaaggactccgccgtgagcaagggcgaggaggataacatggccatcatcaaggagttcatgcgcttcaag<br>gtgcacatggagggctccgtgaacggccacgagttcgagatcgagggcgagggcgagggccgcccctacgagggcacccagaccgccaagctg<br>aaggtgaccaagggtggccccctgcccttcgcctgggacatcctgtcccctcagttcatgtacggctccaaggcctacgtgaagcaccccgcc<br>gacatccccgactacttgaagctgtccttccccgagggcttcaagtgggagcgcgtgatgaacttcgaggacggcggcgtggtgaccgtgacc<br>caggactcctccctgcaggacggcgagttcatctacaaggtgaagctgcgcggcaccaacttcccctccgacggccccgtaatgcagaagaag<br>accatgggctgggaggcctcctccgagcggatgtaccccgaggacggcgccctgaagggcgagatcaagcagaggctgaagctgaaggacggc<br>ggccactacgacgctgaggtcaagaccacctacaaggccaagaagcccgtgcagctgcccggcgcctacaacgtcaacatcaagttggacatc<br>acctcccacaacgaggactacaccatcgtggaacagtacgaacgcgccgagggccgccactccaccggcggcatggacgagctgtacaagggc<br>ggatccggcggaggaagcggccagactctgaaggaaacatcctttaatcaggcctatggtcgagatttaatggaggcccaagagtggtgcagg<br>aagtacatgaaatcagggaatgtcaaggacctcacccaagcctgggacctctattatcatgtgttccgacgaatctcaaagcaggtctccgga<br>ggaggcggcagcggcggagggggatccatggagcatgttgcctttggcagtgaggacatcgagaacactctggccaatatggacgacgaacaa<br>ctggataggttggcctttggcgtaattcagctcgatggtgacgggaatatcctgctgtacaatgctgctgaaggggacatcactggcagagat<br>cccaaacaggtgattgggaagaacttcttcaaggatgttgcacctggaacggatactcccgagttttacggcaaattcaaggaaggcgcagcg<br>tcagggaatctgaacaccatgttcgaatggacgataccgacataa |
| pAG1239 | MYC-pFAST$_{99-114}$-$^{N}$FRB-ECFP-Caax | atggaacaaaagcttatttctgaagaggacttggaattcagcaggggaccaaccaaggtcaaggtgcacttgaagaaagccctttccagcggc<br>gggggaggctccggaggcggaggcagcgagatgtggcatgaaggcctggaagaggcatctcgtttgtactttggggaaaggaacgtgaaaggc<br>atgtttgaggtgctggagcccttgcatgctatgatggaacggggccccggtggtgctagtggtggtgctagtatggtgagcaagggcgaggag<br>ctgttcaccggggtggtgcccatcctggtcgagctggacggcgacgtaaacggccacaagttcagcgtgtccggcgagggcgagggcgatgcc<br>acctacggcaagctgaccctgaagttcatctgcaccaccggcaagctgcccgtgccctggcccaccctcgtgaccaccctgacctggggcgtg<br>cagtgcttcagccgctaccccgaccacatgaagcagcacgacttcttcaagtccgccatgcccgaaggctacgtccaggagcgcaccatcttc<br>ttcaaggacgacggcaactacaagacccgcgccgaggtgaagttcgagggcgacaccctggtgaaccgcatcgagctgaagggcatcgacttc<br>aaggaggacggcaacatcctggggcacaagctggagtacaactacatcagccacaacgtctatatcaccgccgacaagcagaagaacggcatc<br>aaggccaacttcaagatccgccacaacatcgaggacggcagcgtgcagctcgccgaccactaccagcagaacacccccatcggcgacggcccc<br>gtgctgctgcccgacaaccactacctgagcacccagtccgccctgagcaaagaccccaacgagaagcgcgatcacatggtcctgctggagttc<br>gtgaccgccgccgggatcactctcggcatggacgagctgtacaagaagaagaagaagaagaagagcaagaccaagtgcgtgatcatgtaa |
| pAG1224 | MYC-pFAST$_{99-114}$-$^{N}$FRB-ECFP-Cb5 | atggaacaaaagcttatttctgaagaggacttggaattcagcaggggaccaaccaaggtcaaggtgcacttgaagaaagccctttccagcggc<br>gggggaggctccggaggcggaggcagcgagatgtggcatgaaggcctggaagaggcatctcgtttgtactttggggaaaggaacgtgaaaggc<br>atgtttgaggtgctggagcccttgcatgctatgatggaacggggccccggtggtgctagtggtggtgctagtatggtgagcaagggcgaggag<br>ctgttcaccggggtggtgcccatcctggtcgagctggacggcgacgtaaacggccacaagttcagcgtgtccggcgagggcgagggcgatgcc<br>acctacggcaagctgaccctgaagttcatctgcaccaccggcaagctgcccgtgccctggcccaccctcgtgaccaccctgacctggggcgtg<br>cagtgcttcagccgctaccccgaccacatgaagcagcacgacttcttcaagtccgccatgcccgaaggctacgtccaggagcgcaccatcttc<br>ttcaaggacgacggcaactacaagacccgcgccgaggtgaagttcgagggcgacaccctggtgaaccgcatcgagctgaagggcatcgacttc<br>aaggaggacggcaacatcctggggcacaagctggagtacaactacatcagccacaacgtctatatcaccgccgacaagcagaagaacggcatc<br>aaggccaacttcaagatccgccacaacatcgaggacggcagcgtgcagctcgccgaccactaccagcagaacacccccatcggcgacggcccc<br>gtgctgctgcccgacaaccactacctgagcacccagtccgccctgagcaaagaccccaacgagaagcgcgatcacatggtcctgctggagttc<br>gtgaccgccgccgggatcactctcggcatggacgagctgtacaagtccggactcagatctatcaccaccgtggagtccaactcctcctggtgg<br>accaactgggtgatccccgccatctccgccctggtggtggccctgatgtaccgcctatacatggccgaggactag |
| pAG1240 | TOM20-mCherry-$^{C}$FRB-pFAST$_{1-98}$ | atggtgggtcggaacagcgccatcgccgcgggcgtgtgcggtgccctcttcatagggtactgcatctactttgaccgcaaaagacgaagtgac<br>cccaacttcggatccggaggctccggaggaggcgtgagcaagggcgaggaggataacatggccatcatcaaggagttcatgcgcttcaaggtg<br>cacatggagggctccgtgaacggccacgagttcgagatcgagggcgagggcgagggccgcccctacgagggcacccagaccgccaagctgaag<br>gtgaccaagggtggccccctgcccttcgcctgggacatcctgtcccctcagttcatgtacggctccaaggcctacgtgaagcaccccgccgac<br>atccccgactacttgaagctgtccttccccgagggcttcaagtgggagcgcgtgatgaacttcgaggacggcggcgtggtgaccgtgacccag<br>gactcctccctgcaggacggcgagttcatctacaaggtgaagctgcgcggcaccaacttcccctccgacggccccgtaatgcagaagaagacc<br>atgggctgggaggcctcctccgagcggatgtaccccgaggacggcgccctgaagggcgagatcaagcagaggctgaagctgaaggacggcggc<br>cactacgacgctgaggtcaagaccacctacaaggccaagaagcccgtgcagctgcccggcgcctacaacgtcaacatcaagttggacatcacc<br>tcccacaacgaggactacaccatcgtggaacagtacgaacgcgccgagggccgccactccaccggcggcatggacgagctgtacaagggcgga<br>tccggcggaggaagcggccagactctgaaggaaacatcctttaatcaggcctatggtcgagatttaatggaggcccaagagtggtgcaggaag<br>tacatgaaatcagggaatgtcaaggacctcacccaagcctgggacctctattatcatgtgttccgacgaatctcaaagcaggtctccggagga<br>ggcggcagcggcggagggggatccatggagcatgttgcctttggcagtgaggacatcgagaacactctggccaatatggacgacgaacaactg<br>gataggttggcctttggcgtaattcagctcgatggtgacgggaatatcctgctgtacaatgctgctgaaggggacatcactggcagagatccc<br>aaacaggtgattgggaagaacttcttcaaggatgttgcacctggaacggatactcccgagttttacggcaaattcaaggaaggcgcagcgtca<br>gggaatctgaacaccatgttcgaatggacgataccgacataa |
| pAG1225 bis | TOM20-FKBP-pFAST$_{115-125}$ | atggtgggtcggaacagcgccatcgccgcgggcgtgtgcggtgccctcttcatagggtactgcatctactttgaccgcaaaagacgaagtgac<br>cccaacttcggatccagtgctggtggtagtggagtgcaggtggaaaccatctccccaggagacgggcgcaccttccccaagcgcggccagacc<br>tgcgtggtgcactacaccgggatgcttgaagatggaaagaaatttgattcctcccgggacagaaacaagccctttaagtttatgctaggcaag<br>caggaggtgatccgaggctgggaagaaggggttgcccagatgagtgtgggtcagagagccaaactgactatatctccagattatgcctatggt<br>gccactgggcacccaggcatcatcccaccacatgccactctcgtcttcgatgtggagcttctaaaactggaagaatccggaggaggcggcagc<br>ggcggagggggatccggtgacagatattgggtctttgtgaaacgggtgtaa |

| | | |
|---|---|---|
| pAG1258 | MYC -bFos-pFAST$_{115-125}$-IRES-HA-iRFP670 | ATGGAACAAAAGCTTATTTCTGAAGAGGACTTGgaattcggtcgtgcgcagtccatcggtcgtcgcggtaaagttgaacaactgtccccg<br>gaagaggaagagaaacgtcgcatccgccgtgaacgtaacaaaatggcggcagcgaaatgccgtaaccgccgtcgtgaactgaccgacacc<br>ctgcaggcggaaaccgaccagctggaagacgaaaaatccgcgctgcaaaccgaaatcgcgaacctgctgaaagaaaaagaaaagctggag<br>ttcatcctggcggcacaccgtccggcgtgcaaaatcccgaacgacctgggttccggaggaggcggcagcggcggaggggggatccGGTGAC<br>AGATATTGGGTCTTTGTGAAACGGGTGTAACTCGAGGACTACAAGGACGACGACGACAAGCCCGGGatccgcccctctccctcccccccc<br>cctaacgttactggccgaagccgcttggaataaggccggtgtgcgtttgtctatatgttattttccaccatattgccgtcttttggcaat<br>gtgagggcccggaaacctggccctgtcttcttgacgagcattcctaggggtctttcccctctcgccaaaggaatgcaaggtctgttgaat<br>gtcgtgaaggaagcagttcctctggaagcttcttgaagacaaacaacgtctgtagcgaccctttgcaggcagcggaaccccccacctggc<br>gacaggtgcctctgcggccaaaagccacgtgtataagatacacctgcaaaggcggcacaaccccagtgccacgttgtgagttggatagtt<br>gtggaaagagtcaaatggctctcctcaagcgtattcaacaaggggctgaaggatgcccagaaggtaccccattgtatgggatctgatctg<br>gggcctcggtgcacatgctttacatgtgtttagtcgaggttaaaaaaacgtctaggccccccgaaccacggggacgtggttttcctttga<br>aaaacacgatgataatatggCCACaaccATGcgatcgTACCCATACGATGTTCCAGATTACGCTgaattcatggcgcgtaaggtcgatct<br>cacctcctgcgatcgcgagccgatccacatccccggcagcattcagccgtgcggctgcctgctagcctgcgacgcgcaggcggtgcggat<br>cacgcgcattacggaaaatgccggcgcgttctttggacgcgaaactccgcgggtcggtgagctactcgccgattacttcggcgagaccga<br>agcccatgcgctgcgcaacgcactggcgcagtcttccgatccaaagcgaccggcgctgatcttcggttggcgcgacggcctgaccggccg<br>caccttcgacatctcactgcatcgccatgacggtacatcgatcatcgagttcgagcctgcggcggccgaacaggccgacaatccgctgcg<br>gctgacgcggcagatcatcgcgcgcaccaaagaactgaagtcgctcgaagagatggccgcacgggtgccgcgctatctgcaggcgatgct<br>cggctatcaccgcgtgatgttgtaccgcttcgcggacgacggctccgggatggtgatcggcgaggcgaagcgcagcgacctcgagagctt<br>tctcggtcagcactttccggcgtcgctggtcccgcagcaggcgcggctactgtacttgaagaacgcgatccgcgtggtctcggattcgcg<br>cggcatcagcagccggatcgtgcccgagcacgacgcctccggcgccgcgctcgatctgtcgttcgcgcacctgcgcagcatctcgccctg<br>ccatctcgaatttctgcggaacatgggcgtcagcgcctcgatgtcgctgtcgatcatcattgacggcacgctatggggattgatcatctg<br>tcatcattacgagccgcgtgccgtgccgatggcgcagcgcgtcgcggccgaaatgttcgccgacttcttatcgctgcacttcaccgccgc<br>ccaccaccaacgc |
| pAG1265 | MYC -bFos(Δ179-193)-pFAST$_{115-125}$-IRES-HA-iRFP670 | ATGGAACAAAAGCTTATTTCTGAAGAGGACTTGgaattcggtcgtgcgcagtccatcggtcgccgcggtaaagttgaacaactgtccccg<br>gaagaggaagagaaacgtcgcatccgccgtgaacgtaacaaaatggcggcagcgaaatgccgtaaccgccgtcgtgaactgaccgacacc<br>ctgcaggcggaaaccgaccagctggaagacgaaaaatccgcggagttcatcctggcggcacaccgtccggcgtgcaaaatcccgaacgac<br>ctgggttccggaggaggcggcagcggcggaggggggatccGGTGACAGATATTGGGTCTTTGTGAAACGGGTGTAACTCGAGGACTACAAG<br>GACGACGACGACAAGCCCGGGatccgcccctctccctcccccccccctaacgttactggccgaagccgcttggaataaggccggtgtgcg<br>tttgtctatatgttattttccaccatattgccgtcttttggcaatgtgagggcccggaaacctggccctgtcttcttgacgagcattcct<br>aggggtctttcccctctcgccaaaggaatgcaaggtctgttgaatgtcgtgaaggaagcagttcctctggaagcttcttgaagacaaaca<br>acgtctgtagcgaccctttgcaggcagcggaaccccccacctggcgacaggtgcctctgcggccaaaagccacgtgtataagatacacct<br>gcaaaggcggcacaaccccagtgccacgttgtgagttggatagttgtggaaagagtcaaatggctctcctcaagcgtattcaacaagggg<br>ctgaaggatgcccagaaggtaccccattgtatgggatctgatctggggcctcggtgcacatgctttacatgtgtttagtcgaggttaaaa<br>aaacgtctaggccccccgaaccacggggacgtggttttcctttgaaaaacacgatgataatatggCCACaaccATGcgatcgTACCCATA<br>CGATGTTCCAGATTACGCTgaattcatggcgcgtaaggtcgatctcacctcctgcgatcgcgagccgatccacatccccggcagcattca<br>gccgtgcggctgcctgctagcctgcgacgcgcaggcggtgcggatcacgcgcattacggaaaatgccggcgcgttctttggacgcgaaac<br>tccgcgggtcggtgagctactcgccgattacttcggcgagaccgaagcccatgcgctgcgcaacgcactggcgcagtcttccgatccaaa<br>gcgaccggcgctgatcttcggttggcgcgacggcctgaccggccgcaccttcgacatctcactgcatcgccatgacggtacatcgatcat<br>cgagttcgagcctgcggcggccgaacaggccgacaatccgctgcggctgacgcggcagatcatcgcgcgcaccaaagaactgaagtcgct<br>cgaagagatggccgcacgggtgccgcgctatctgcaggcgatgctcggctatcaccgcgtgatgttgtaccgcttcgcggacgacggctc<br>cgggatggtgatcggcgaggcgaagcgcagcgacctcgagagctttctcggtcagcactttccggcgtcgctggtcccgcagcaggcgcg<br>gctactgtacttgaagaacgcgatccgcgtggtctcggattcgcgcggcatcagcagccggatcgtgcccgagcacgacgcctccggcgc<br>cgcgctcgatctgtcgttcgcgcacctgcgcagcatctcgccctgccatctcgaatttctgcggaacatgggcgtcagcgcctcgatgtc<br>gctgtcgatcatcattgacggcacgctatggggattgatcatctgtcatcattacgagccgcgtgccgtgccgatggcgcagcgcgtcgc<br>ggccgaaatgttcgccgacttcttatcgctgcacttcaccgccgcccaccaccaacgc |
| pAG1713 | H2B-pFAST$_{99-114}$-P2A-bJUN- pFAST$_{1-98}$-IRES-HA-mTurquoise2 | ATGCCCGAACCTGCGAAGTCAGCGCCCGCTCCCAAAAAAGGCTCTAAAAAAGCTGTCGCCAAGACCCAGAAGAAGGGGGATAAGAAAAGG<br>CGTAAGACCAGGAAAGAGAGTTACGCCATTTACGTGTACAAAGTACTAAAACAAGTCCACCCGGACACTGGCATCTCCTCAAAGGCGATG<br>GGCATTATGAACTCATTTGTAAACGACATCTTCGAGCGCATCGCCGGAGAAGCGTCGCGCCTGGCGCATTACAACAAGCGCTCCACTATC<br>ACATCCCGGGAGATCCAGACGGCCGTGCGCCTGCTCCTGCCCGGAGAACTGGCCAAACACGCTGTGTCTGAGGGCACAAAGGCCGTGACC<br>AAGTACACCAGCTCCAAGAGCAGGGGACCAACCAAGGTCAAGGTGCACTTGAAGAAAGCCCTTTCCagcggcgggggaggctccggaggc<br>ggaggcagcGGAAGCGGAGCTACTAACTTCAGCCTGCTGAAGCAGGCTGGAGACGTGGAGGAGAACCCTGGACCTAaggcggagaggaag<br>cgcatgagaaaccgcatcgctgcctccaagtgccggaaaaggaagctggagcggatcgcccggctagaggaaaaagtgaaaaccttgaaa<br>gcgcaaaactccgagctggcgtccacggccaacatgctcagggaacaggtggcacagcttaaacagaaagtcatgaaccacgttaacagt<br>gggtgccaactcatgctaacgcagcagttgcaaacgtttTccggaggaggcggcagcggcggaggggggatccATGGAGCATGTTGCCTTT<br>GGCAGTGAGGACATCGAGAACACTCTGGCCAATATGGACGACGAACAACTGGATAGGTTGGCCTTTGGCGTAATTCAGCTCGATGGTGAC<br>GGGAATATCCTGCTGTACAATGCTGCTGAAGGGGACATCACTGGCAGAGATCCCAAACAGGTGATTGGGAAGAACTTCTTCAAGGATGTT<br>GCACCTGGAACGGATACTCCCGAGTTTTACGGCAAATTCAAGGAAGGCGCAGCGTCAGGGAATCTGAACACCATGTTCGAATGGACGATA<br>CCGACATAACTCGAGGACTACAAGGACGACGACGACAAGCCCGGGatccgcccctctccctcccccccccctaacgttactggccgaagc<br>cgcttggaataaggccggtgtgcgtttgtctatatgttattttccaccatattgccgtcttttggcaatgtgagggcccggaaacctggc<br>cctgtcttcttgacgagcattcctaggggtctttcccctctcgccaaaggaatgcaaggtctgttgaatgtcgtgaaggaagcagttcct<br>ctggaagcttcttgaagacaaacaacgtctgtagcgaccctttgcaggcagcggaaccccccacctggcgacaggtgcctctgcggccaa<br>aagccacgtgtataagatacacctgcaaaggcggcacaaccccagtgccacgttgtgagttggatagttgtggaaagagtcaaatggctc<br>tcctcaagcgtattcaacaaggggctgaaggatgcccagaaggtaccccattgtatgggatctgatctggggcctcggtgcacatgcttt<br>acatgtgtttagtcgaggttaaaaaaacgtctaggccccccgaaccacggggacgtggttttcctttgaaaaacacgatgataatatggC<br>CACaaccATGcgatcgTACCCATACGATGTTCCAGATTACGCTgaattcatggtgagcaagggcgaggagctgttcaccggggtggtgcc<br>catcctggtcgagctggacggcgacgtaaacggccacaagttcagcgtgtccggcgagggcgagggcgatgccacctacggcaagctgac<br>cctgaagttcatctgcaccaccggcaagctgcccgtgccctggcccaccctcgtgaccaccctgtcctggggcgtgcagtgcttcgcccg<br>ctaccccgaccacatgaagcagcacgacttcttcaagtccgccatgcccgaaggctacgtccaggagcgcaccatcttcttcaaggacga<br>cggcaactacaagacccgcgccgaggtgaagttcgagggcgacaccctggtgaaccgcatcgagctgaagggcatcgacttcaaggagga<br>cggcaacatcctggggcacaagctggagtacaactacttcagcgacaacgtctatatcaccgccgacaagcagaagaacggcatcaaggc<br>caacttcaagatccgccacaacatcgaggacggcggcgtgcagctcgccgaccactaccagcagaacacccccatcggcgacggccccgt<br>gctgctgcccgacaaccactacctgagcacccagtccaagctgagcaaagaccccaacgagaagcgcgatcacatggtcctgctggagtt<br>cgtgaccgccgccgggatcactctcggcatggacgagctgtacaagtaa |
